# Monitoring fish communities through environmental DNA metabarcoding in the fish pass system of the second largest hydropower plant in the world

**DOI:** 10.1101/2021.08.17.456687

**Authors:** Giorgi Dal Pont, Camila Duarte Ritter, Andre Olivotto Agostinis, Paula Valeska Stica, Aline Horodesky, Nathieli Cozer, Eduardo Balsanelli, Otto Samuel Mäder Netto, Caroline Henn, Antonio Ostrensky, Marcio Roberto Pie

## Abstract

The Itaipu Hydroelectric Power Plant is the second largest in the world in power generation. The artificial barrier created by its dam imposes an obstacle for fish migration. Thus, in 2002, a fish pass system, named Piracema Channel, was built to allow fish to access areas upstream of the reservoir. We tested the potential of environmental DNA metabarcoding to monitor the impact of both the dam and associated fish pass system in the Paraná River fish communities and to compare it with traditional monitoring methods. Using a fragment of the 12S gene, we characterized richness and community composition based on amplicon sequence variants, operational taxonomic units, and zero-radius OTUs. We combined GenBank and in-house data for taxonomic assignment. We found that different bioinformatics approaches showed similar results. Also, we found a decrease in fish diversity from 2019 to 2020 probably due to the recent extreme drought experienced in southeastern Brazil. The highest alpha diversity was recorded in the mouth of the fish pass system, located in a protected valley with the highest environmental heterogeneity. Despite the clear indication that the reference databases need to be continuously improved, our results demonstrate the analytical efficiency of the metabarcoding to monitor fish species.

## Background

The Itaipu Hydroelectric Power Plant, built at the border between Brazil and Paraguay, is the second largest in the world in power generation^1^, second only to the Three Gorges Power Plant in China. With the formation and filling of its reservoir, in 1982^2^, the natural barrier to the migration of fishes of the middle section of the Paraná River (Sete Quedas falls) was replaced by the artificial barrier of the Itaipu dam, located 170 km downstream. This artificial barrier (196 m high) caused impacts on the adjacent fish assemblages, such as the reduction in reproductive activity in the first kilometers downstream of the dam^3^. To allow for fish migration and mitigate the environmental impact of the dam, a fish passage system known as the Piracema Channel was created in 2002, linking the Paraná River to Itaipu’s Reservoir^4^. However, the real contribution to the reproductive success of the long-distance migratory species is still under investigation, and this channel also allowed for the dispersal of species originally restricted to the lower Paraná River upstream and species originally restricted to the upper Paraná River downstream^5^. These potential impacts are continuous and can interact with natural disturbance, such as several droughts as which happened in 2020. In this context, monitoring the impact of both dam and fish pass system in the Paraná River fish communities is essential.

Fish diversity estimates in Brazilian freshwater are still imprecise due to the scarcity of complete inventories^5–7^. Many species are described every year and several groups are in need of taxonomic revision^5,8^. Furthermore, traditional assessment methods for fish diversity surveys are costly and time consuming, given that they depend on capture (e.g. netting, trawling) or observation^9,10^ and expertise for taxonomic identification^11^. In this sense, designing methods for cost-effective monitoring fish diversity and community composition is an urgent task. Most sampling efforts in Brazil have historically been primarily funded by the hydroelectric sector, focusing particularly on rivers where power dams were built^12^. The areas of the dam construction have some of the most comprehensive knowledge of fish assemblage composition in comparison with other Brazilian regions and therefore offer an ideal opportunity to compare taxonomic surveys with molecular approaches.

A promising alternative to traditional taxonomic surveys and biomonitoring methods is the use of environmental DNA (eDNA), combined with a high-throughput sequencing approach, as in the case of metabarcoding^13^. This technique has the advantage of obtaining DNA from environmental samples, such as water, without first isolating the target organism and therefore can sample entire communities^14^. Metabarcoding is a powerful tool for biodiversity assessment that has been widely used for several purposes and different taxonomic groups^15–17^, and is considered a transformative technology for the entire field^18^. However, some limitations, such as the relative scarcity of DNA sequences for several species, which is even more problematic in highly diverse regions such as the Neotropics^19^, may create constraints that hamper its full application^20,21^.

The absence of a comprehensive DNA reference database may lead to a misidentification of several species. Therefore, putting together a curated and complete DNA reference database is fundamental for species identification through a metabarcoding approach^7^. But, even with an incomplete DNA reference database, the use of molecular units, such taxonomic units clustered by similarity (operational taxonomic units - OTUs^22^) or unique sequences (e.g. amplicon sequence variants - ASVs^23^, or zero-radius OTUs - ZOTUs^24^) allows for diversity monitoring in the context of biodiversity assessment in megadiverse biomes. Such estimates without comprehensive species identification limit ecological conclusions but allowed for monitoring of natural and artificial impacts^16,25^. The metabarcoding approach has been successfully used for molecular identification of several vertebrate groups in temperate regions^26,27^, monitoring of endangered species such as freshwater fish in Australia^28^ and turtles in the United States^29^, and to describe biodiversity even with limited taxonomic identification^30^.

In this context, our goal here is to describe an effective survey protocol for detecting fish assemblages through eDNA metabarcoding in an ecologically complex and highly diverse freshwater system, the Piracema channel, that connects the Paraná River with Itaipu Reservoir. For this, we used an in-house molecular database of fishes occurring in the channel complemented with GenBank sequences. We also describe fish alpha diversity and community structure in the Piracema channel system. Additionally, we compare our metabarcoding results with the traditional sampling campaigns made between 2017-2021.

## Material and Methods

### Study area

Our study was conducted at the Piracema channel (Fig. 1), a fish pass system connecting the Paraná River with the Upper Paraná River floodplain (main reservoir). For the traditional taxonomic survey, we sampled three points at mouth of channel at Paraná River (Fig. 1, blue square), the main lake at the Piracema channel (Fig. 1, red circle), and the reservoir near the water intake to the Channel (Fig. 1, green triangle) between 2017 to 2021. Fish were collected monthly during the fish reproductive period (October to March), and once during the winter (July or August), employing active and passive methods (Table 1).

**Figure 1.**
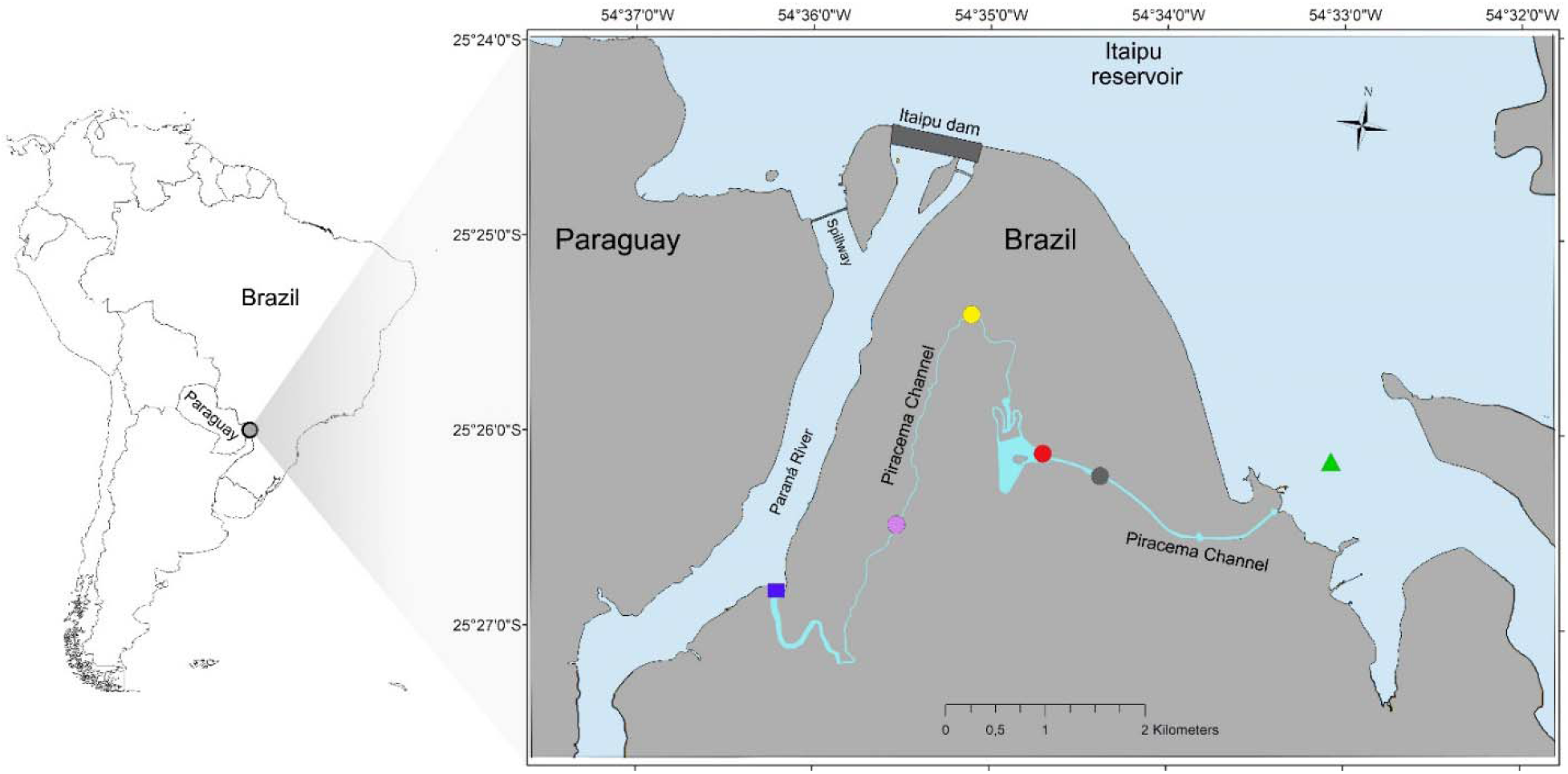
Sampling location. The map shows the sampling location of each collection point. We sampled one point at mouth of channel at Paraná River (blue square), four points along the Piracema Channel (circles; Bela Vista River 1 = purple, Bela Vista 2 = yellow, Brasilia stream = gray, and lake = red), and one point at Itaipu’s reservoir (green triangle). Up at figure is possible to visualize the Itaipus’ dam that created the reservoir. Inset panel shows the location of Itaipu’s dam in relation to South America. Map was created in QGIS v.3.6.2 software ^88^.

**Table 1.** Fish sampling methods at the Piracema Channel

| **Fish**<br>**sampling**<br>**method** | **Quantitative and qualitative aspects** |
| --- | --- |
| Gill nets | Mesh sizes: 1 a 10 cm (adjacent knots), each one 10m long and 1 to 1,5m high |
| Longlines | 30 hooks, 10 of each size: /10, /8 and /6, fish pieces as bait |
| Cast nets | Mesh sizes: 3 e 6 cm (adjacent knots) |
| Electrofishing | Smith-Root, backpack electrofisher, 600V, 30Hz DC |

For each point, gill nets and longlines were set out in the afternoon (16:00 h) and inspected every 4 hours during a 24 h cycle; cast nets were operated 3 times each mesh, after every gear inspection. Boarded electrofishing was operated two times in each point, at dawn and at dusk, covering 100m of the environment margin each time. Fish were euthanized by immersion in benzocaine solution, following current legislation^31^, and identified accordingly Britski et al.^32^, Ota et al.^33^ and Neris et al.^34^. Fragments of muscle were collected with a scalpel, placed into 2 ml tubes filled with 99.8% ethanol and stored at 4ºC until processing. Voucher specimens are housed in the Nupelia-UEM fish collection.

For metabarcoding, we sampled one site at the mouth of channel at Paraná River (Fig. 1, blue square), four sites along the Piracema Channel (Fig. 1, circles), and one site at Itaipu Reservoir (Fig. 1, green triangle). Each sampling point was collected in sextuplicate. All six sites were sampled in 2019 and three sites (mouth of channel at Paraná River, lake at Piracema channel [red circle], and the reservoir) were sampled again in 2020, totaling 54 samples. All sampling sites were provided with GPS coordinates.

### Sampling design for molecular analysis

We collected water by partially submerging a one litter polypropylene bottle. The objective was to sample water at the air/water interface. After water collection, bottles were closed and cleaned with a 10% sodium hypochlorite solution, following by rinsing with distilled water. We used gloves which were changed in between each new sampling replicate to reduce the risk of cross-sample contamination.

After the collection and cleaning steps, the bottles were stored in polystyrene boxes containing artificial ice to maintain the temperature of the samples at 4 to 10 °C. The samples were filtered, on the same day of collection, using nitrocellulose membranes (0.45 μm pore) with the aid of a vacuum pump. Filters were kept in 100% ethanol under refrigerated conditions until molecular analysis was performed. All filters were processed at the ATGC laboratory at the Universidade Federal do Paraná (UFPR).

### DNA extraction

For total DNA extraction, we kept the collected filters at room temperature to allow the residual ethanol to dry completely. After dried we extract the DNA using magnetic beads (microspheres surrounded by magnetite and carboxyl), which bind to DNA (carboxyl bond - DNA) by the process of Solid Phase Reversible Immobilization (SPRI). The DNA extract was stored at –20 °C until the amplification. The extraction and quantification processes were carried out in separate rooms, as suggested by Pie et al.^35^. We checked the DNA concentration using both a spectroscope (Nanodrop, Thermo, USA) and a fluorimeter (Qubit, Invitrogen, USA).

### PCR amplification

We targeted the 12S rRNA gene using the MiFish forward (5’-GTCGGTAAAACTCGTGCCAGC-3’) and reverse (5’-CATAGTGGGGTATCTAATCCCAGTTTG-3’) primers designed by Miya et al.^36^ to yield 163– 185 bases long fragments. Amplification was performed in a total volume of 20 μl in GoTaqG2 system (Promega, USA), 500 nM of forward and reverse primers, and 20 ng of DNA template. The PCR conditions consisted of an initial denaturation step of 2 min at 95 °C and then 25 cycles of denaturation at 94 °C for 30 s, hybridization at 55 °C for 45 s, and elongation at 72 °C for 30 s, followed by a final elongation at 72 °C for 5 min and finishing at 4 °C. To avoid PCR inhibition BSA (0.5 μg/μl) was added to the reaction as suggested by Boeger et al^37^. The quality of amplification was verified on a 1.5% agarose gel in TBE buffer (9 mM TRIS, 9 mM boric acid, 1 mM EDTA), stained with SYBR Safe DNA Gel Stain (ThermoFisher Scientific, Country). All replicates from each sampling point were amplified to increase the chance of detecting rare species. The PCR product was then diluted (20x) and used as a template for the addition of adapters in the second PCR. Indexing was performed for Illumina MiSeq sequencing (Illumina, USA), using the above PCR system with Nextera indexes (Illumina) in a total volume of 10 μl. PCR conditions were an initial step of 95 °C for 3 min, following by 12 cycles of denaturation at 94 °C for 30 s, hybridization at 55 °C for 45 s, and elongation at 72 °C for 30 s, followed by a final elongation at 72 °C for 5 min and finishing at 4 °C. We checked the DNA concentration in a Qubit^®^ fluorimeter (Invitrogen, USA), normalized and pooled the PCR products following the Illumina protocol. The samples were sequenced at GoGenetic (Curitiba, Brazil) using Illumina MiSeq Reagent 600V3 (PE 300b). Three negative controls (distilled water) were used as control for extraction, amplification, and sequencing. The raw sequences are deposited in GenBank under Bioproject PRJNA750895 (biosamples SAMN20500524 – SAMN20500577).

### Sequence analyses and taxonomic assessment

For the amplicon sequence variants (ASVs) approach, we used the Cutadapt package^38^ in Python v.3.3^39^ to remove primers. We then used the DADA2 package^23^ in R v. 4.0.2^40^ to quality filter reads, merge sequences, remove chimeras, and to infer ASVs. We excluded reads with ambiguous bases (maxN=0). Based on the quality scores of the forward and reverse sequences, each read was required to have <3 or <5 errors, respectively (maxEE=c (3,5), truncQ=2). Therefore, ASVs were inferred for forward and reverse reads for each sample using the run-specific error rates. To assemble paired-end reads, we considered a minimum of 12 base pairs of overlap and excluded reads with mismatches in the overlapping region. Chimeras were removed using the consensus method of “removeBimeraDenovo” implemented in DADA2.

For operational taxonomic units (OTUs) and zero-radios OTU (ZOTUs) analyses, we used the USEARCH/UPARSE v.11.0.667 Illumina paired reads pipeline^41^ to primer remove, quality filtering, dereplicate and sort reads by abundance, to infer OTUs and ZOTUs, and to remove singletons. We filtered the sequences to discard chimeras and clustered sequences into OTUs at a minimum similarity of 97% using a ‘greedy’ algorithm that performs chimera filtering and OTU clustering simultaneously and the UNOISE algorithm to denoised sequence as zero-radios OTUs to create or ZOTUs table^41,42^.

We build a reference dataset of DNA sequences for the 205 fish taxa that have been historically recorded in the Itaipu system using the following steps. First, we looked for 12S sequences of these species in GenBank by searching for their corresponding names. We were able to find sequences for 126 species in our reference database. Additionally, we created an in-house database which included sequences for 42 additional species to the 79 species previously identified as present on Itaipu system but not available on GenBank. Sequences for the in-house database were obtained via Sanger sequencing of tissue samples and were uploaded to GenBank (accession numbers MZ778813-MZ778856). We manually blasted all sequences against the NCBI GenBank database to verify misidentification or problematic sequences (e.g. blasted in the different family). In total, our reference database included 168 (82%) sequences from the 205 taxa recorded in the Itaipu system. Finally, we blasted the ASVs, OTUs, and ZOTUs sequences with our reference database to verify the taxonomic composition using the “Blastn” function of the program Blast+^43^ with an e-value of 0.001. We kept ASVs, OTUs, and ZOTUs that have matched with a fish species at minimum level of 75% similarity (as these sequences are probably fishes species not present in our reference database), and considered identified species just ASVs, OTUs, and ZOTUs that matched in a minimum level of 97% similarity. We summed reads for each ASVs, ZOTUs, and OTUs present in the three negative controls and divided by the total reads of each ASVs, ZOTUs, and OTUs. All ASVs, ZOTUs and OTUs with a proportion > 0.01% of reads in negative controls were discarded (13 ASVs, 2 ZOTU, and 7 OTUs).

### Statistical analysis

We conducted all analyses in R using RStudio^44^. We used the tidyverse package v. 1.3.0^45^ for data curation and ggplot2 v. 3.3.2^46^, ggfortify v. 0.4.11^47^, gridExtra v. 2.3^48^, and ggpubr v. 0.4.0^49^ for data visualization (scripts in Appendix 1).

For analysis of alpha and beta diversity with metabarcoding data, we made the analysis at ASVs, OTUs, and ZOTUs level. Since the number of observed ASVs, ZOTUs, and OTUs is dependent on the number of reads, we rarefied all samples to the lowest number of reads obtained from any one plot (157 for ASVs, 147 for ZOTUs, and 219 for OTUs; Fig. S1) using the “rarefy” function with the vegan v.2.5.7^50^ R package. Because in the ZOTUs table the minimum reads of a plot was nine, we used the second lower value to avoid having to downsize the other samples to such a low number of reads^51^. Because rarefying of counts is considered inappropriate for detection of differentially abundant species^51^, even more with so different sampling depth as in our case, we also calculated true effective number of ASVs, ZOTUs, and OTUs of order *q*=1, which is equivalent to the exponential of the Shannon entropy^52^, using the function “AlphaDiversity” of the Entropart v.1.6.7^53^ R package. The effective number is more robust against biases arising from uneven sampling depth than the simple counts of ASVs, ZOTUs, and OTUs^51,54^. Additionally, for alpha diversity, we also calculated the ASV, OTU, and ZOTU richness (the number of ASV, OTU, and ZOTU per point), Chao1, and Fisher’s alpha diversity (i.e., the relationship between the number of ASV, OTU, and ZOTU in any given point and the number of reads of each ASV, OTU, and ZOTU) using the phyloseq v.1.34.0^55^ R package.

For beta diversity, we also used rarefaction (with “rrarefy” function of vegan package) and hill number (with “varianceStabilizingTransformation” function in DESeq2 v.1.28.1^56^ R package) to normalize our data. While rarefaction normalizes data by random subsampling without replacement, the hill number transformation normalizes the count data with respect to sample size (number of reads in each sample) and variances, based on fitted dispersion-mean relationships^56^. We then constructed two-dimensional Principal Coordination Analysis (PCoA) ordinations of the abundance (reads) and presence/absence data for both rarefied and hill numbered data. We used the ‘cmdscale’ function and Bray-Curtis distances in the vegan package to assess community dissimilarity among all samples in the PCoA. We used the “envfit” method in vegan to fit sampling localities and sample year onto the PCoA ordination as a measure of the correlation among the sampling localities with the PCoA axes.

For traditional survey data, we calculated the alpha diversity using the observed richness, Chao1, and ACE with the function “estimate”, and Shannon index with the function “diversity” both with vegan package. We also constructed two-dimensional PCoA ordinations of the abundance (reads) and presence/absence data, and used the “envfit” to fit sampling localities and sample year onto the PCoA ordination.

## Results

For the traditional surveys, 4,447 fishes were collected, for a total 122 species. Most specimens were collected at the mouth of channel at Paraná River, with 2,240 (51%) fishes belonging to 105 species. The reservoir showed the lowest number of collected specimens: 1,034 (23%) and a total of 64 species. Of these, 29 species (24%) were identified with metabarcoding approach, while 93 species (76%) were only identified with traditional surveys (Table 2). Other 27 species were identified at > 97% of similarity only with the metabarcoding approach (Table 2).

**Table 2.**
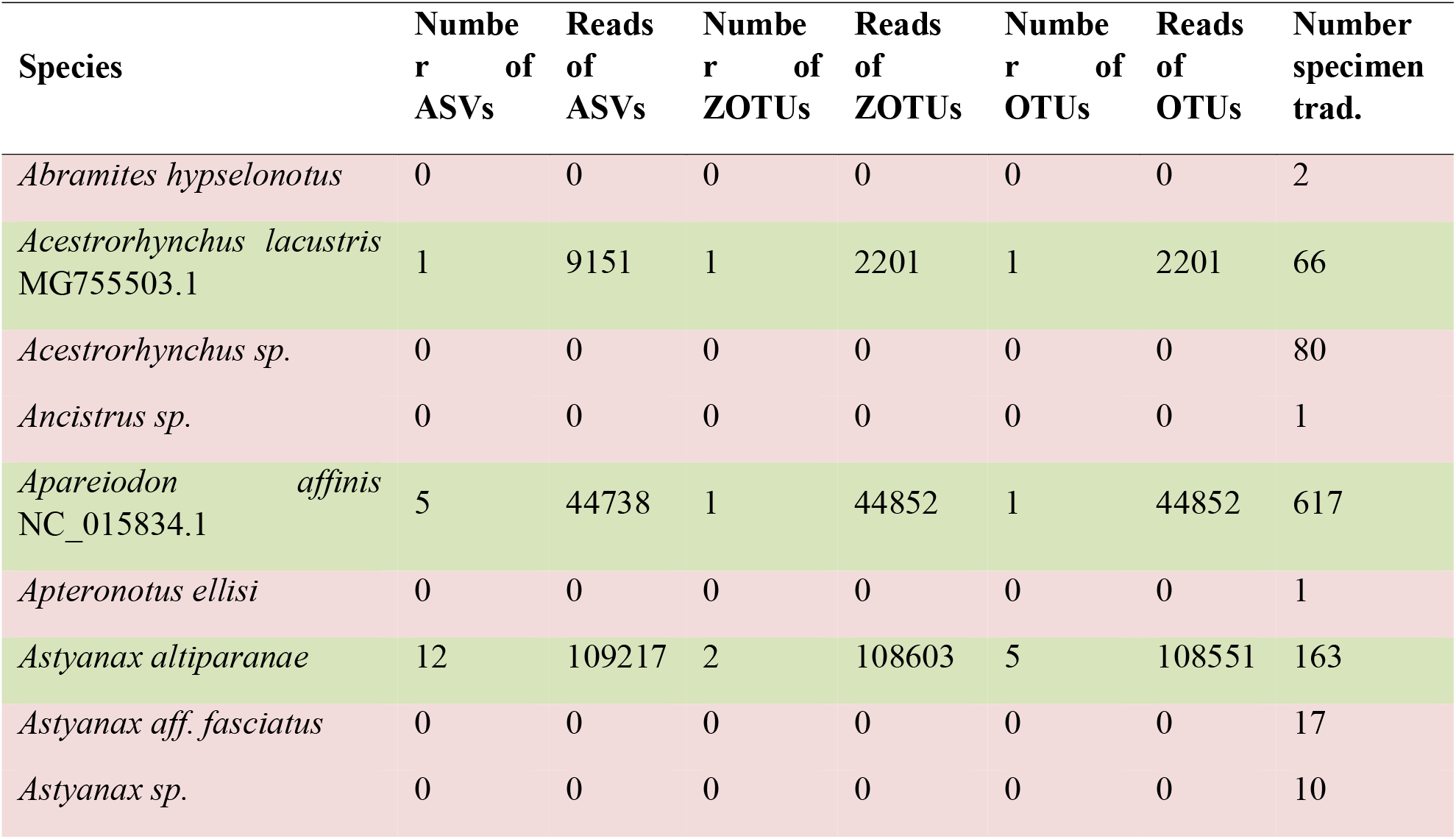
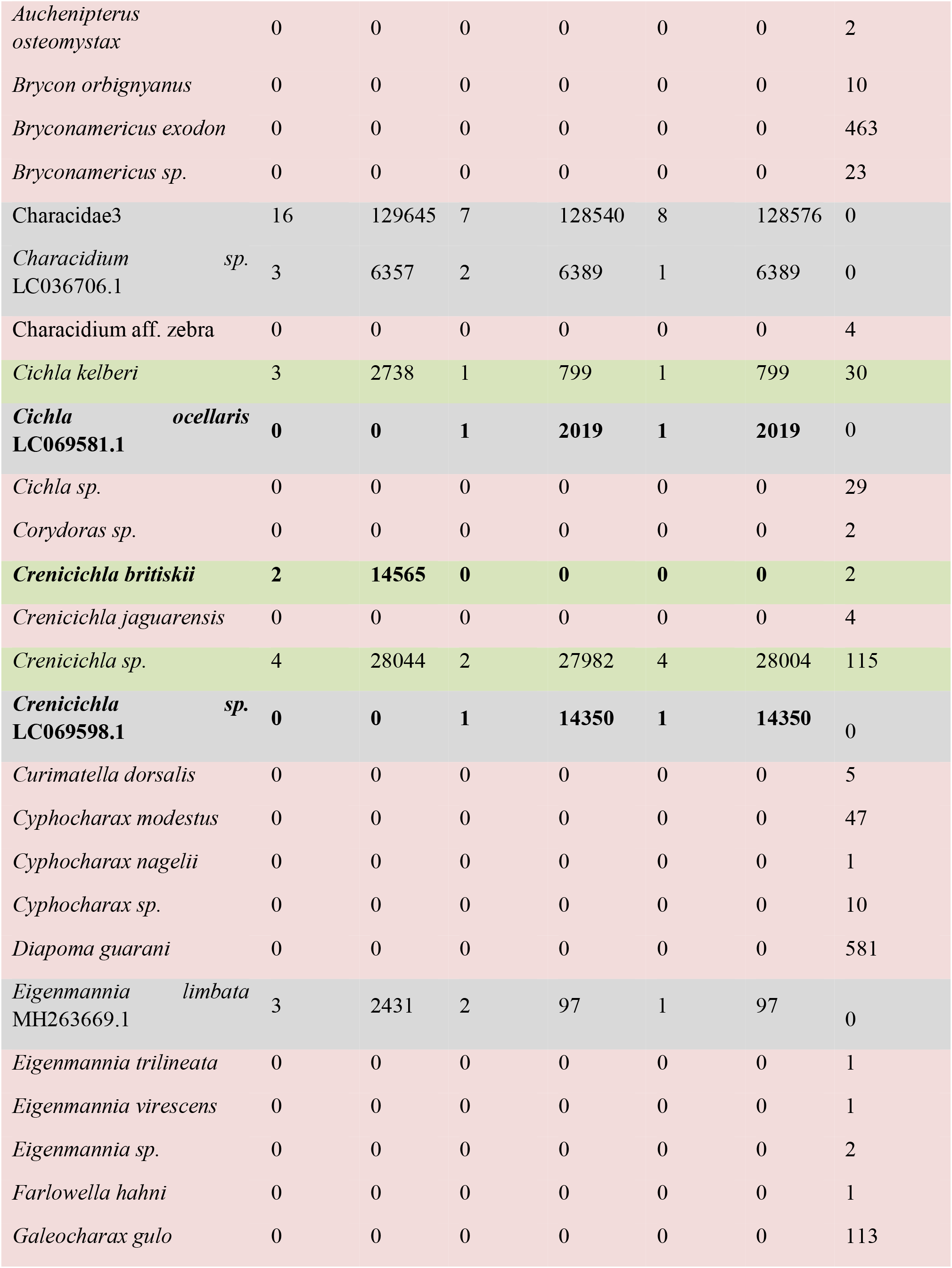
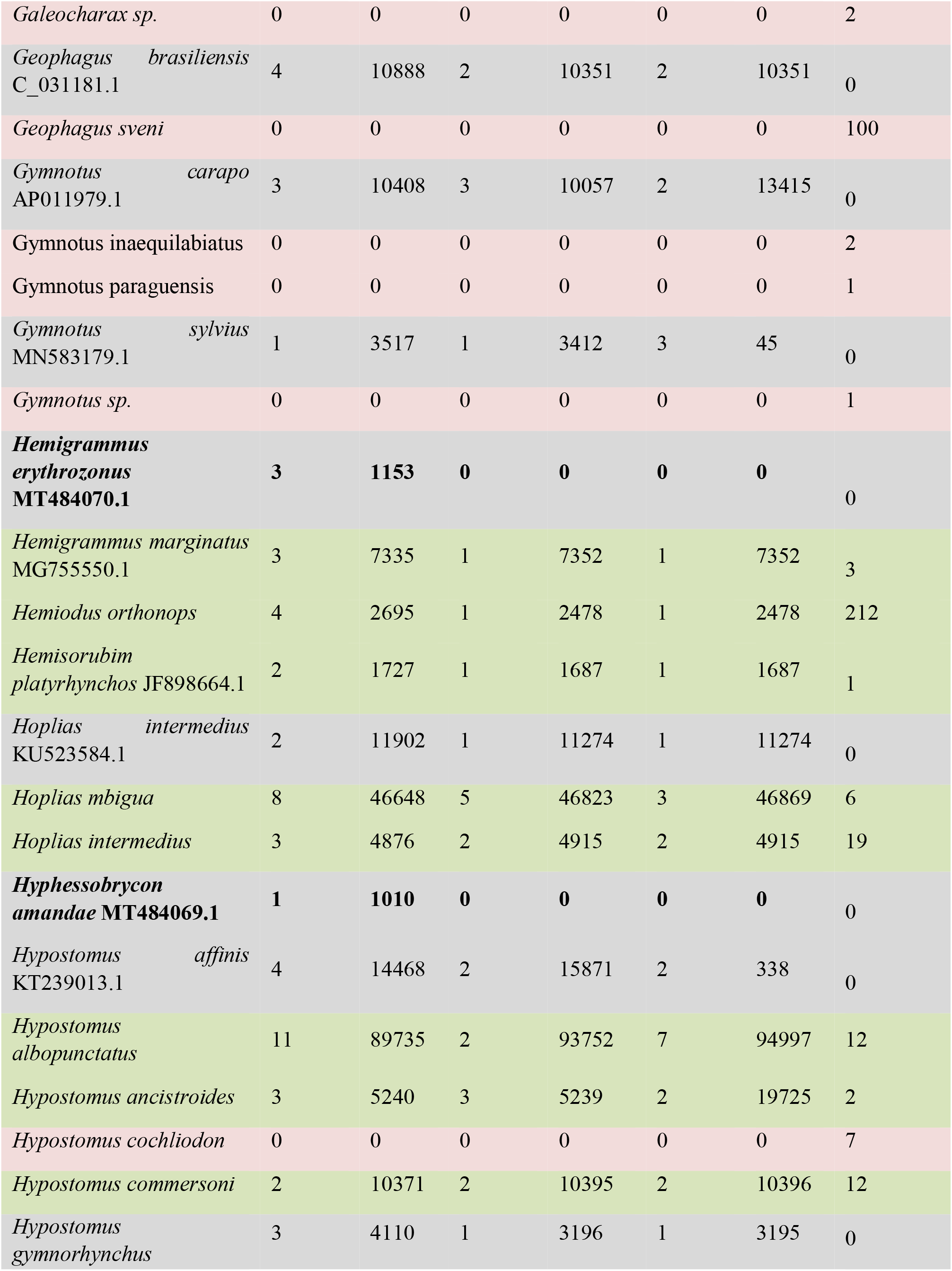
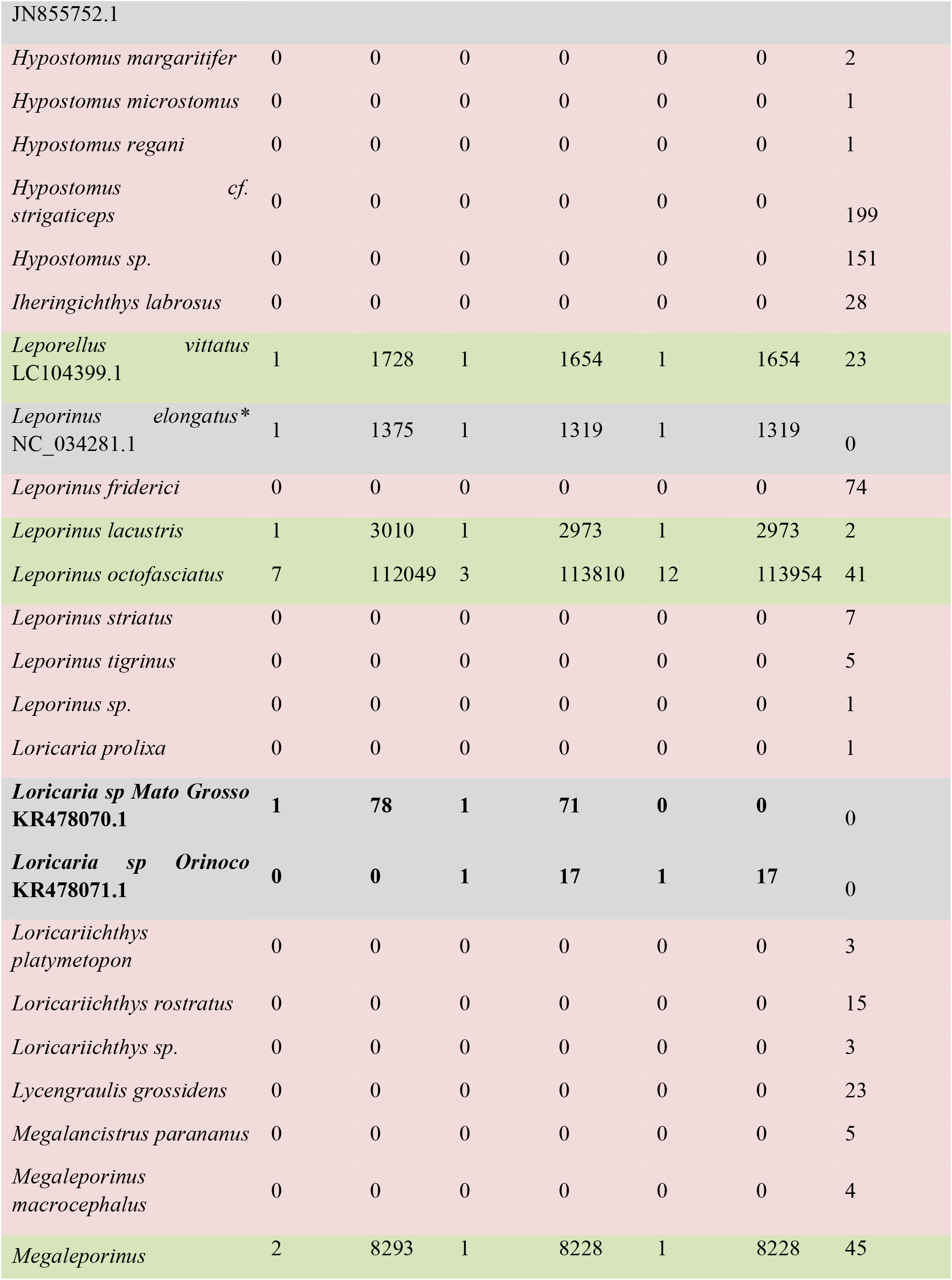
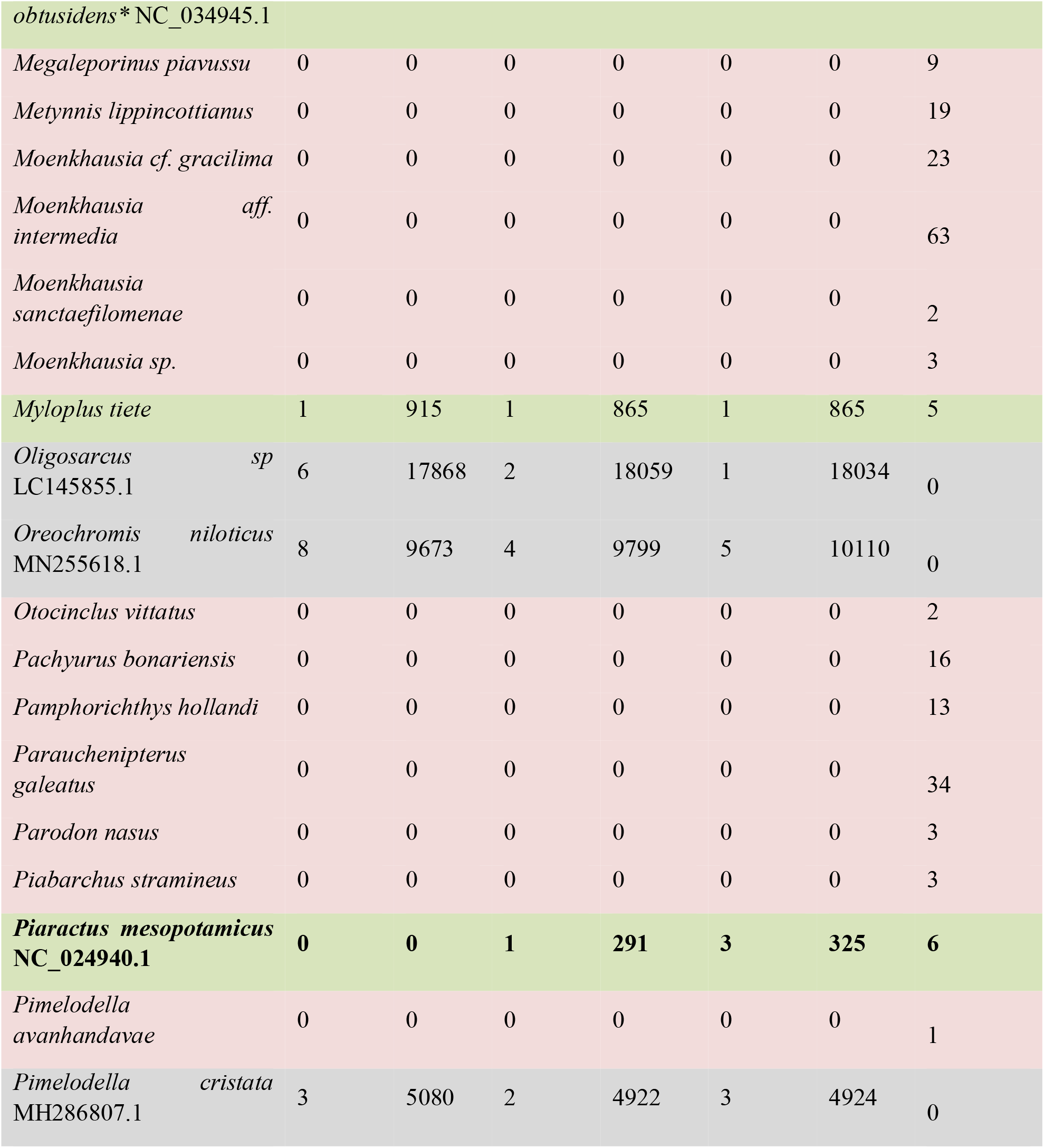
Number of specimens sampled with traditional surveys (N. specimen trad.), species identified at level of > 97% similarity, the number of ASVs, ZOTUs, OTUs identified per species (possible intra-specific variation) and the number of reads per species after correction. Rows in green are species identified in both traditional surveys and metabarcoding (29 in total), red are species just identified by traditional surveys (93 in total), and gray species just identified by metabarcoding approach (27 in total). In bold species that was not registered in one of the pipelines. *Leporinus elongatus* is now *Megaleporinus obtusidens*, but as both species names are in GenBank and different ASVs, ZOTUs, and OTUs match each sequence, we keep both and marked with an asterisk (*).

For metabarcoding data, we obtained a total of 17,616,032 reads. After all cleaning steps, we kept a total of 2,280,447sequences belonging to 7,096 ASVs. Of these, 1,015,157 (44% of the total) sequences belonging to 190 ASVs were classified into species corresponding to our reference database in the level of 75% similarity. Of these, 7,591 reads (0.75% of the total) were recorded in the sum of the three negative controls (Appendix 3). A total of 13 ASVs with a proportion of reads > 0.01% of the total reads were removed from the analysis. A total of 121 ASVs (64%) were classified in 35 species matches at a level > 97% of similarity (Table 2), which is certainly an underestimation of the real number of species, since the other 69 ASVs should belong to species do not present in our database.

For the OTU and ZOTU analyses, after all cleaning steps, we obtained 1,157,738 and 1,145,129 reads belonging to 796 OTUs and 207 ZOTUs, respectively. Of these, 1,002,883 (87%) and 1,002,335 (87%) reads belonging to 136 OTUs and 94 ZOTUs, respectively, were classified into species corresponding to our reference database at the level of 75% similarity. Of these, 7,493 and 7,494 reads (0.75% of total) were registered in the sum of the three negative controls in the OTUs and ZOTUs tables, respectively (Appendix 4 for OTUs and 5 for ZOTUs) and 2 ZOTU and 7 OTUs with a proportion of reads > 0.01% of the total reads were removed from the analysis. As the OTUs analysis already classified the sequences by 97% similarity, the 131 OTUs (all matched with fishes less the 5 present in the negative controls) probably correspond to the number of species present in our samples (more than all species sampled in five years with traditional surveys). Yet, only 37 species belonging to 42 OTUs and 34 species belonging to 46 ZOTUs at > 97% similarity were assigned in both analyses. Eighty-one (66%) OTUs and 41 (47%) ZOTUs were identified as a fish species with a similarity lower than 97%, representing species not present in our reference database.

All the alpha diversity measures of ASVs, OTUs, and ZOTUs per sampling point varied, with the point at mouth of the channel at Paraná River, in 2019, showing the highest diversity for all molecular units and the lake at Piracema Channel in 2020 the lowest (Fig. 2). For the traditional surveys, the variation was more random, but the mouth of channel at Paraná River also had the highest diversity (Fig. S2).

**Figure 2.**
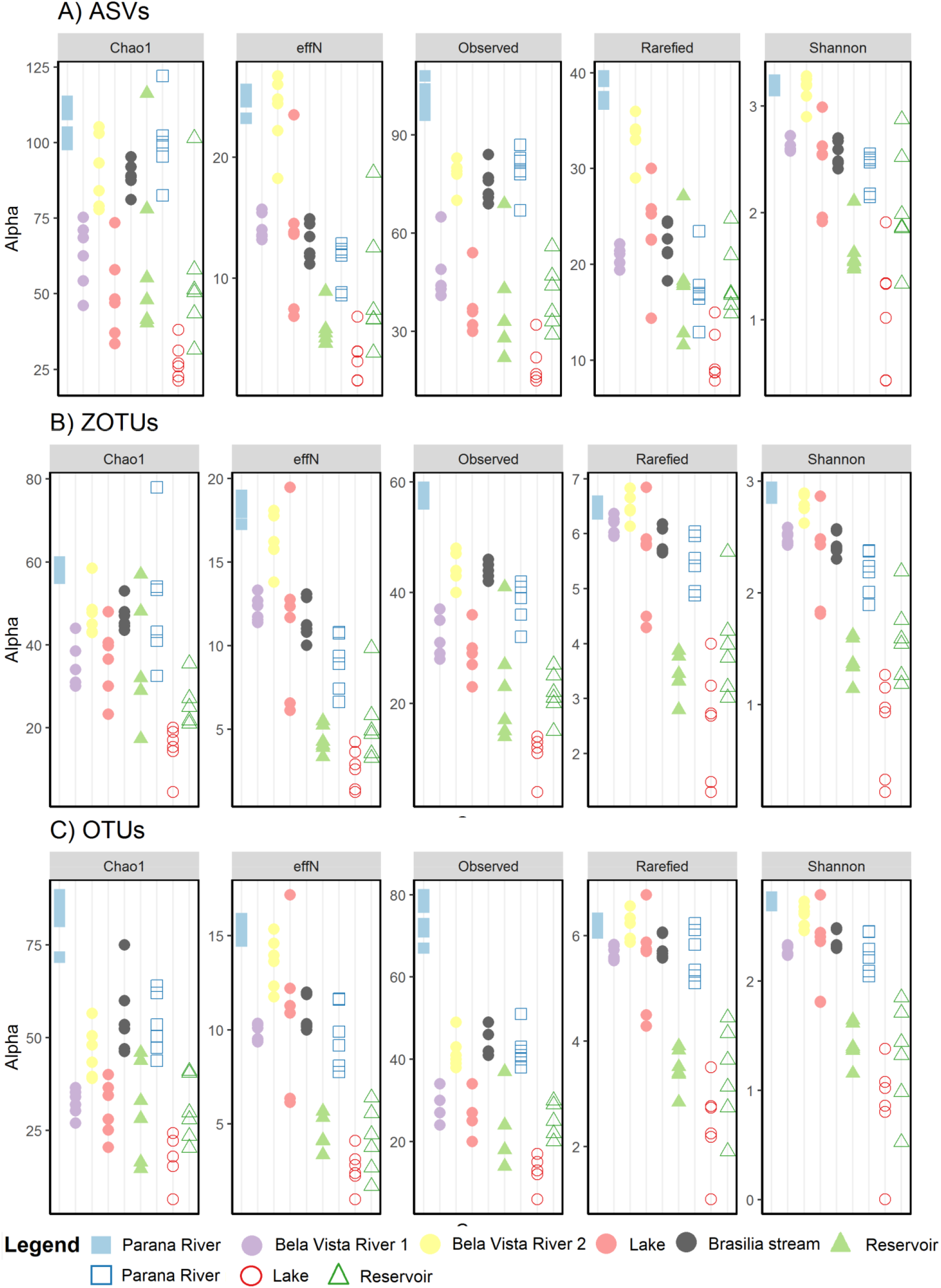
Alpha diversity estimation for A) ASVs, B) ZOTUs, C) OTUS. Alpha diversity varied by location and by sampling year. Each point is one of the replicates sampled. Colors and symbols represent collection points (mouth of channel at Paraná River = blue square, Itaipu’s reservoir = green triangle, and Piracema Channel = circles [Bela Vista River 1 = purple, Bela Vista 2 = yellow, Brasilia stream = gray, and lake = red]), and fill represent year of collection (filled = 2019, empty = 2020).

Fish communities varied among sampled sites. For the abundance based in hill numbers, the first axis of the PCoA separated the samples by year (envfit: R^2^ = 0.30 [ZOTUs], 0.34 [ASVs], and 0.36 [OTUs], p < 0.001), except for the Itaipu’s reservoir, with the positive values associated with 2019 and the negative values associated with 2020 (Fig. 3). The second axis separated the samples by locality with some overlap (envfit: R^2^ = 0.95 [ZOTUs] - 0.96 [ASVs and OTUs], p < 0.001; Fig. 3). For the presence/absence data also based in hill numbers, the overlap was higher but yet the separation by year (envfit: R^2^ = 0.30[ZOTUs], 0.32 [OTUs], and 0.41 [ASVs], p < 0.001) and locality (envfit: R^2^ = 0.91 [ZOTUs] - 0.92 [ASVs and OTUs], p < 0.001) was similar to the abundance data (Fig.3). The results of rarefied data were similar with more overlap among sampling points (Appendix 1, Fig. S3). For traditional surveys, the points are clustered by localities (envfit: R^2^ = 0.60 for abundance and 0.87 for presence/absence data, p < 0.001), but not by year (p >0.05, Fig. S4).

**Figure 3.**
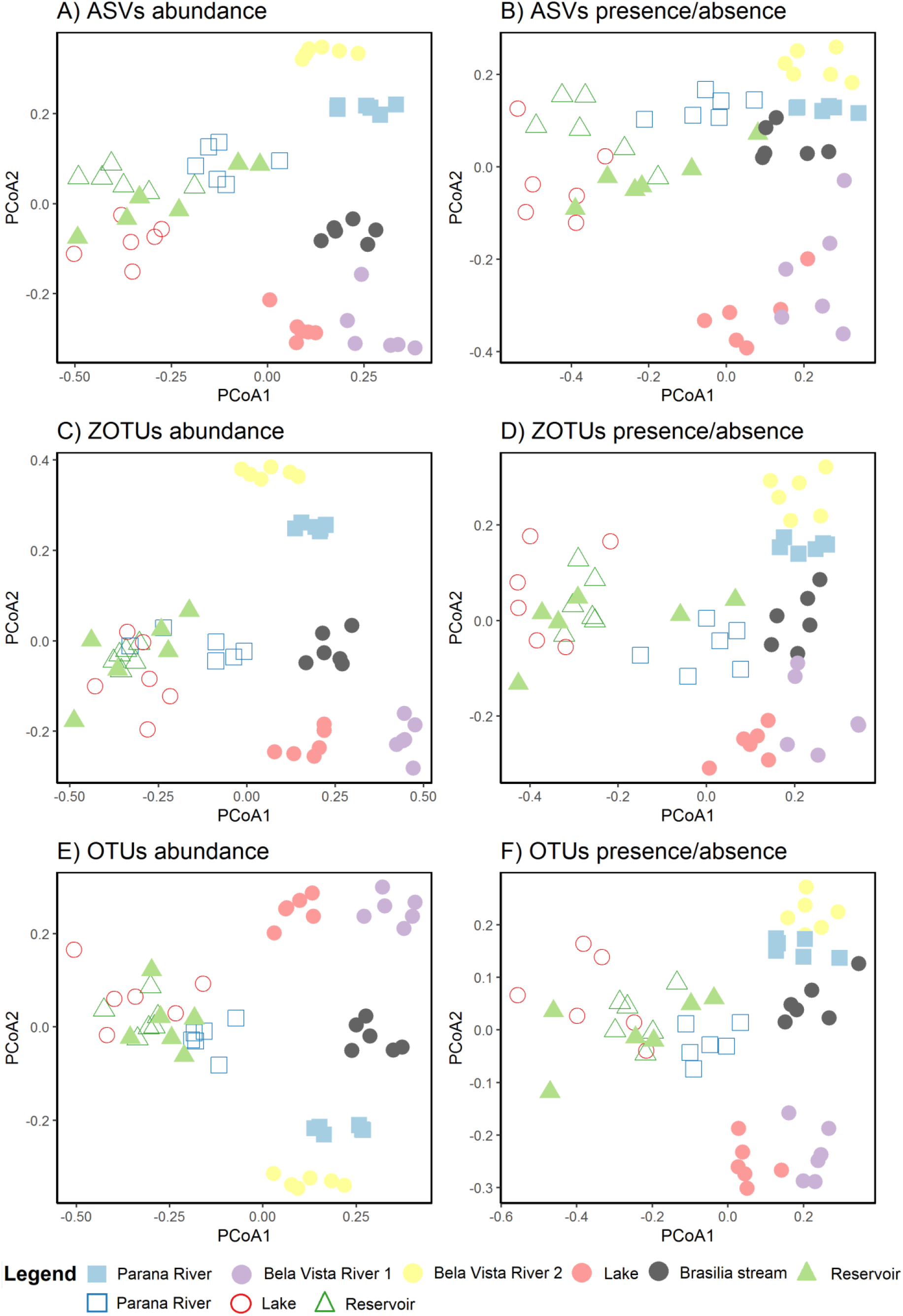
Principal Coordinates Analysis (PCoA) of fishes’ communities from Itaipu based in hill numbers for A) ASVs abundance, B) ASVs presence/absence, C) ZOTUs abundance, D) ZOTUs presence/absence, E) OTUs abundance, and F) OTUs presence/absence. The axis 1 separated mainly the samples by year, while the axis 2 separated samples mainly by locality. Each point is one of the replicates sampled. Colors and symbols represent collection points (mouth of channel at Paraná River = blue square, Itaipu’s reservoir = green triangle, and Piracema Channel = circles [Bela Vista River 1 = purple, Bela Vista 2 = yellow, Brasilia stream = gray, and lake = red]), and filled represent year of collection (fill = 2019, empty = 2020).

## Discussion

Our results support mounting evidence that eDNA analysis provides a cost-effective alternative to characterize fish biodiversity. We also demonstrate that different bioinformatic approaches show similar results in terms of alpha and beta diversity, supporting the use of molecular approaches to monitor biodiversity even with incomplete taxonomic identification. However, a serious caveat for using these molecular methods for biodiversity assessments is the scarcity of comprehensive taxonomic reference databases, especially for the tropical regions of the globe. Here, we also highlight these caveats for the Neotropical fish database, which are taxonomically limited, limiting the identification of several species. With a complete reference database, eDNA could detect mostly fish community and also fish species that are poorly or non-represented by conventional methods, as suggested by our results.

We identified 35 species with ASVs, 37 with OTUs, and 34 with ZOTUs approaches at >97% similarity. However, many other ASVs, OTUs, and ZOTUs were identified at <97% similarity, representing species not present in our database. Considering that 76 species sampled with traditional survey had no available sequences, many of these species may be present in our metabarcoding data but blasted as another species. We produced our reference database based on the historical taxonomic survey of Piracema Channel that may prevent identification of species that had not been recorded by conventional fish survey methods. However, the use of a database without curatorship can include spurious species identifications, such as species unlikely to be physically present at sampling sites^10^. That occurs because when the database does not contain the sequence of a certain species, the sequences will match with the closest species in that database, which can occur in a completely different environment (e.g. marine), beyond other factors that also contribute to registering spurious species, such as misannotated sequences^57^ or low variability in the target sequenced region^10^ that will sign any species with such similar sequence. For instance, our sequence for *Prochilodus lineatus* is identical to other *Prochilodus* species, such as *P. harttii* and *P. costatus*. Furthermore, there are many species undescribed, making it impossible to identify them. A recent compilation to list Paraná state fish species included 42 undescribed species^5^, and this number may be underestimated due to the presence of crypt species and sampling biases.

Even with the previously mentioned limitations, the use of molecular units such as ASVs^58^, OTUs^22^, and ZOTUs^24^ allows for assessing of genetic diversity and enables comparison among multiple sites^59^, space-time dynamics^16^ and evaluate natural and anthropogenic impacts^60^. For instance, vertebrate populations from freshwater ecosystems are declining at alarming rates (83% decline since 1970)^61^, and their conservation and management are a priority for global biodiversity^62^. The Neotropical region harbors one of the largest freshwater biodiversity, with an estimated 9,000 described fish species (around 30% of total freshwater species)^11^. The increasing construction of dams is threatening fish populations over the entire planet ^63–65^ but specially in Neotropical countries such as Brazil^5,66,67^, and effective ways to monitor fish biodiversity to understand its impact is essential.

As observed with the use of conventional ichthyofauna monitoring methods^68^, the number of species, ASVs, OTUs, ZOTUs, or 12S gene sequence readouts identified in our study showed a variation between the two sampling occasions (2019 and 2020). Such variations in fish assemblages can be related to a series of factors, both biotic (ecological characteristics of the species, for example) and abiotic (variations in water quality, and other environmental factors). In addition, physical characteristics of the environment such as total water volume and hydrological characteristics can also play a key role in the ecology and occurrence of fish species^68^. For instance, the recent extreme drought experienced in southeastern Brazil^69^ may have impacted fish assemblages. Our results showed a decrease of alpha diversity in 2020 in both mouth of channel at Paraná River (blue squares) and the lake (red circles; Fig. 2). In addition to the direct effects caused by this type of climatic phenomenon, such as the reduction in the volume of water, indirect effects such as reduced oxygen concentration in the water and food availability can cause severe impacts on fish’s communities^68,70,71^. Such effects were more evident at the mouth of channel at Paraná River, where the water level dropped 7 m from 2019 to 2020. At the reservoir, alpha diversity did not vary as water level fluctuation was less evident as a result of a stable environment due to the large size of this water body (green triangles; Fig. 2). However, the traditional survey in Piracema Channel was unable to significantly detect the diversity variation throughout the period of the study (Fig. S2), highlighting the high sensibility of eDNA metabarcoding for monitoring.

Among sampling points, the highest alpha diversity was recorded in those collected in mouth of channel at Paraná River, while the lowest alpha diversity was registered in the lake (Fig. 2). Habitat heterogeneity is recognized as a main factor supporting functional and phylogenetic diversity, which is often reflected in the taxonomic richness of the fish communities^72^. Mouth of channel at Paraná River, the entrance of the Piracema Channel, is in a protected valley, where the riparian vegetation is conserved, allowing the colonization by a diversified flora and fauna. Besides this, the confluence with the Paraná River produces adjacent lotic and lentic microhabitats, supporting a higher alpha diversity when compared to the main lake or the water intake of the Channel, which are lentic and uniform environments. Such pattern of fish diversity / limnologic gradients meets the patterns previously assessed for the reservoir tributaries^73^.

The beta diversity showed that in 2020, with the event of the extreme drought, a homogenization of fish assemblage happened (Fig. 3). Both samples from the mouth of channel at Paraná River (blue squares) and the lake (red circles) cluster together with the reservoir in both years. The Itaipu’s Reservoir was filled in 1982 and the Piracema Channel (a fish pass), connecting the region just downstream from Itaipu Dam to the Itaipu Reservoir, was opened 38 years later. Both events allowed the dispersion of species (including non-native species) in both directions promoting the homogenization of communities from upper and lower Paraná River^5,74,75^. Our results show the importance of the closest rivers and streams for system diversity and resilience, as the mostly community variation was found in the Boa Vista River and Brasilia Stream (Fig. 3).

Although eDNA metabarcoding is a powerful tool for biodiversity, as it has been widely used for different purposes and different taxonomic groups, including identification and quantification of Neotropical ichthyofauna^16,76,77^, many issues can hamper the metabarcoding results^7,10,78,79^. Shaw et al.^10^ drew attention to methodological considerations related to the eDNA sampling process for freshwater fishes. According to them, the number of replicates is extremely important to obtain accurate data. Specifically, they demonstrated that the collection of two eDNA replicates per point were insufficient to detect less abundant taxa; however, adopting five replicates must have a 100% detection rate. In addition, sampling water column was more effective in detecting fish communities than sampling sediment^10^. Here, we collected six replicates per sampling point on the water surface. Furthermore, the rarefaction curves clearly show that many individual samples have a very low sequencing depth, but considering the replicates all our sampled localities reach the asymptote (Fig. S1), although samples from the lake in 2020 just reach the asymptote considering OTUs analyses and the reservoir in 2020 had not reached the asymptote considering ZOTUs analyses (Fig. S1).

The bioinformatic methodological choices can also affect the metabarcoding results. Here, we used three pipelines that showed the best results compared with other approaches^80^. We used both OTU-level clustering at 97% level, with UPARSE^41^, and the unique sequences with zero-radius ZOTU-level denoising, with UNOISE3^24^, and ASV-level Divisive Amplicon Denoising Algorithm 2, with DADA2^23^. Both the OTUs and the ZOTUs are created using in USEARCH^81^. The initial steps as merging, filtering, and deduplicating are the same for both approaches, with just the last step been different. The third approach generated ASVs through a parametric model, based in Q-scores to calculate a substitution model, estimating a probability for each possible base substitution, to infer true biological sequences from reads as implemented in DADA2^23^. Although we recorded some variation in the number of reads and “species” registered in each pipeline, the results are very similar, highlighting their robustness.

Another potential bias in the results is data treatment. Here we used several data normalizations for both alpha and beta diversity. Although historically more used, rarefied data is biased to detect differentially abundant species^51^ and the hill numbers are considered the best approach for metabarcoding data^54^. Also, due to PCR biases, variation in the copy number of 12S genes per cell/genome, as well as differences in size and biomass across the targeted organisms can compromise a straightforward interpretation of OTU reads as an abundance measure^82–84^. However, rare (low abundant) ASVs, ZOTUs and OTUs are more likely to be an artefact (both erroneous sequence or because of cross-talk^85^) and the true sequences are more stochastically distributed due to the intrinsic low occurrence and detection probability^86,87^. Therefore, analyses that weight more the most abundant molecular units could be preferable. As each method has its own biases, we present here both approaches.

Finally, it is important to highlight that, in general, molecular data derived from “environmental sequencing” should be seen as complementary to, rather than as competing with, traditional taxonomic studies. Indeed, a confluence of both lines of evidence is highly warranted, as it will be necessary to overcome their respective shortcomings. For instance, we have shown here that many species occurring in the Itaipu fish pass system have no genetic data to allow their identification. Even so, other 47 species with sequences available were only identified with traditional surveys. This difference can be related to species density but also to primers biases. To improve the species detection with metabarcoding it is crucial to enhance the genetic reference database through traditional and to test if the primers used in metabarcoding studies are able to amplify species present in the studied system. Indeed, the metabarcoding approach is an intricate web of feedback loops with the species taxonomy and ecology.

## Conclusion

Despite the clear indication that the reference databases need to be continuously fed with additional information on species that occur in the region, our results demonstrate the analytical efficiency of the metabarcoding approach for monitoring fish species in the Itaipu’s fish pass system. In addition, the methodology allowed, even when the specific identity of the ASVs, OTUs, and ZOTUs were below 97% similarity with the species in our database, to carry out estimates of species alpha and beta diversity. The use of such a methodology enables the monitoring of the fish community with sufficient sensitivity to detect changes due to some natural or anthropogenic event.

## Acknowledgements

Funding was provided by Itaipu Binacional to AON (grant #4500049847). We thank Itaipu Binacional for providing postdoctoral fellow grants to GDP and NC and the Alexander van Humboldt foundation for providing a postdoctoral fellow grant to CDR. We also thank Conselho Nacional de Desenvolvimento Científico e Tecnológico (CNPq) for awarding AO and MRP with research fellowship grants (grants #304633/2017-8 and #302904/2020-4, respectively).

## Ethics statement

We confirm that all methods were carried out in accordance with relevant guidelines and regulations and in compliance with the ARRIVE guidelines. We confirm that all experimental protocols were in accordance with the precepts of Law nº 11.794, of 8 October 2008, of Decree nº 6.899, of 15 July 2009, and with the edited rules from Conselho Nacional de Controle da Experimentação Animal (CONCEA), and it was approved by the ANIMAL USE ETHICS COMMITTEE OF THE AGRICULTURAL SCIENCES CAMPUS OF THE FEDERAL UNIVERSITY OF PARANA, BRAZIL, with degree 1 of invasiveness, on March 22th, 2021.

## Notes

### Competing Interest Statement

The authors have declared no competing interest.

