## Appendix 2 for "Monitoring fish communities through environmental DNA metabarcoding in the fish pass system of the second largest hydropower plant in the world"

Giorgi Dal Pont^1,2£^, Camila Duarte Ritter^1,2,3*£^, Andre Olivotto Agostinis^1^, Paula Valeska Stica^1,2^, Aline Horodesky^1,2^, Nathieli Cozer^1,2^, Eduardo Balsanelli^4^, Otto Samuel Mäder Netto^2^, Caroline Henn^5^, Antonio Ostrensky^1,2^, Marcio Roberto Pie^1,2^

^1^ Grupo Integrado de Aquicultura e Estudos Ambientais, Departamento de Zootecnia, Universidade Federal do Paraná, Rua dos Funcionários, 1540, Juvevê, 80035-050 Curitiba, PR, Brazil.

^2^ ATGC Genética Ambiental LTDA. Rua dos Funcionários 1540, Juvevê, Curitiba, PR Brazil, 80035-050

^3^ Eukaryotic Microbiology, Faculty of Biology, University of Duisburg-Essen, Universitätsstrasse 5, D-45141 Essen, Germany.

^4^ Departamento de Bioquímica e Biologia Molecular, Universidade Federal do Paraná, Rua dos Funcionários, 1540, Juvevê, 80035-050 Curitiba, PR, Brazil.

^5^ Itaipu Binacional. Divisão de Reservatório - MARR.CD, Avenida Tancredo Neves, 6731, Foz do Iguaçu, Paraná, CEP 85866-900, Brazil

*Corresponding author: Camila D. Ritter,. Phone: +55 48991434597. Postal address: Eukaryotic Microbiology, Faculty of Biology, University of Duisburg-Essen, Universitätsstrasse 5, S05 R04 H83, D-45141 Essen, Germany.

£ Joint first authorship

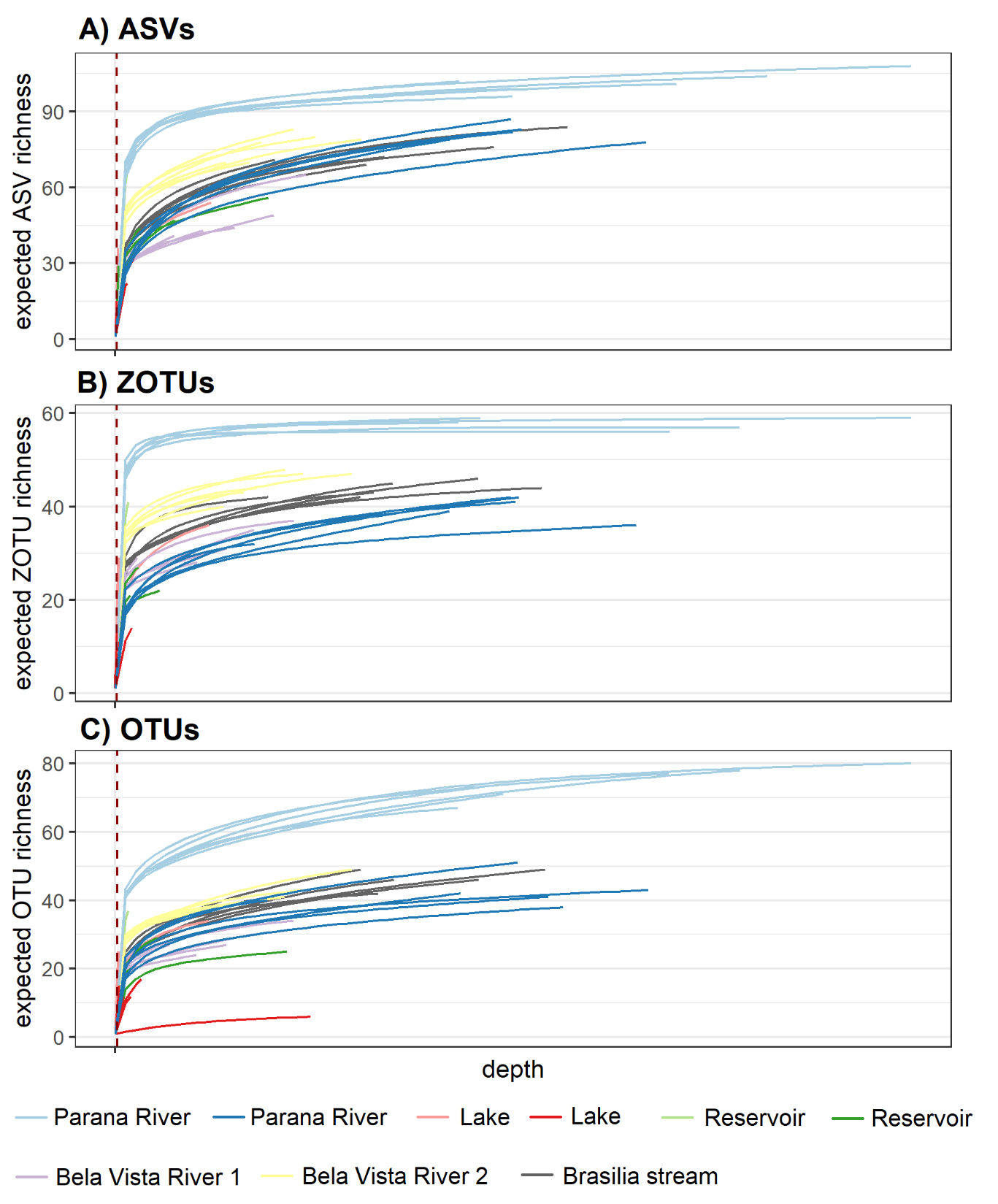

Figure S1**.** Rarefaction curves by sample for the A) ASV, B) ZOTUs, and C) OTUs approaches. The red line shows the smallest number of reads in a single sample, so that all samples were downsized to this number of reads. Colors represent collection points (mouth of Paraná River = blue, Itaipu’s reservoir = green, and Piracema Channel = [Bela Vista River 1 = purple, Bela Vista 2 = yellow, Brasilia stream = gray, and lake = red]), and tone represent year of collection (light = 2019, dark = 2020).

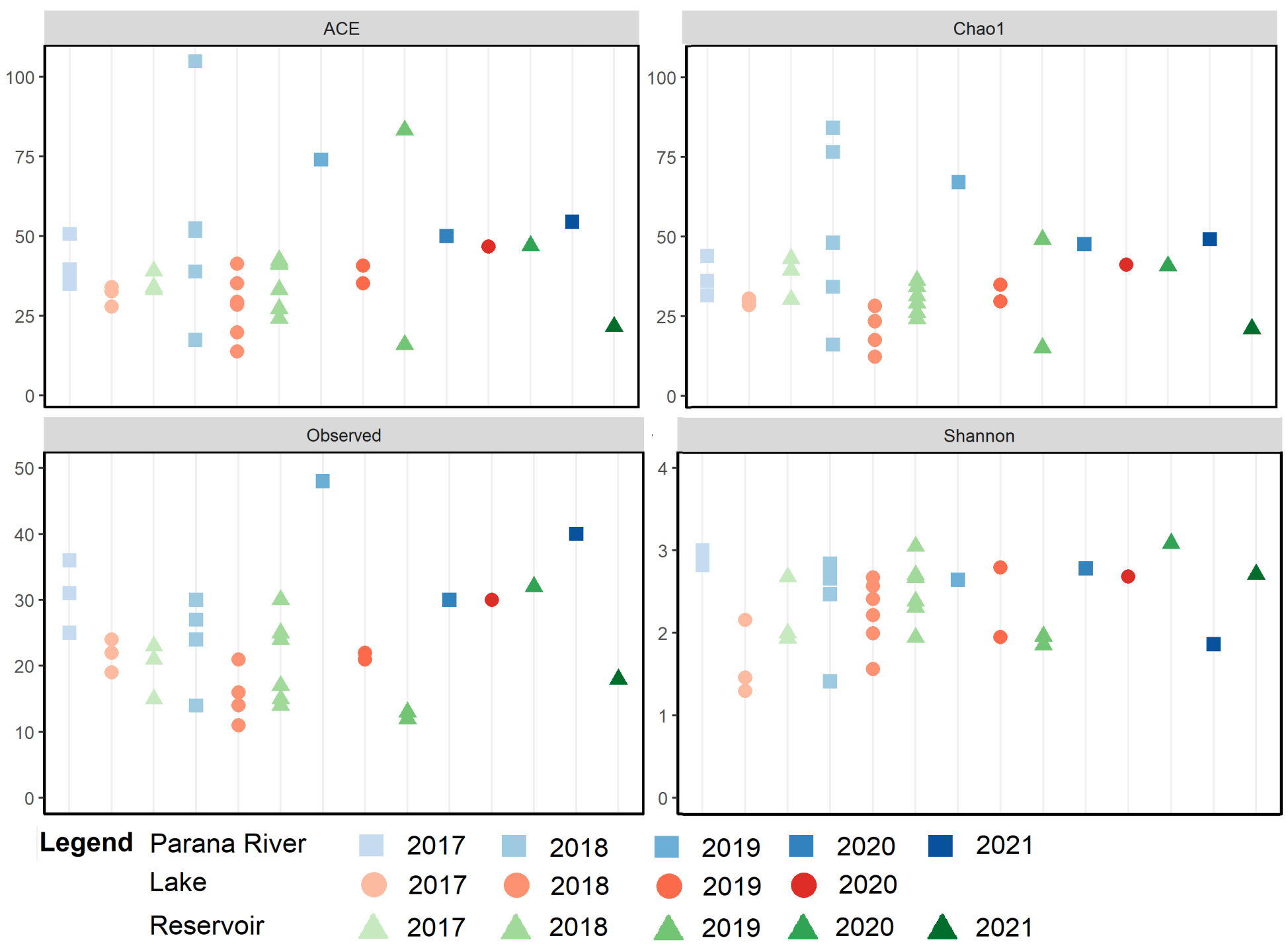

**Figure S2.** Alpha diversity estimation for Itaipu’s fish samples based in the traditional surveys. Alpha diversity varied by location and by sampling year. Each point is one of the sampled replicates. Colors and symbols represent collection sites (mouth of Paraná River = blue square, Itaipu’s reservoir = green triangle, and lake at Piracema Channel = red circles, and tone represent year of collection (lightest = 2017to darkest = 2021).

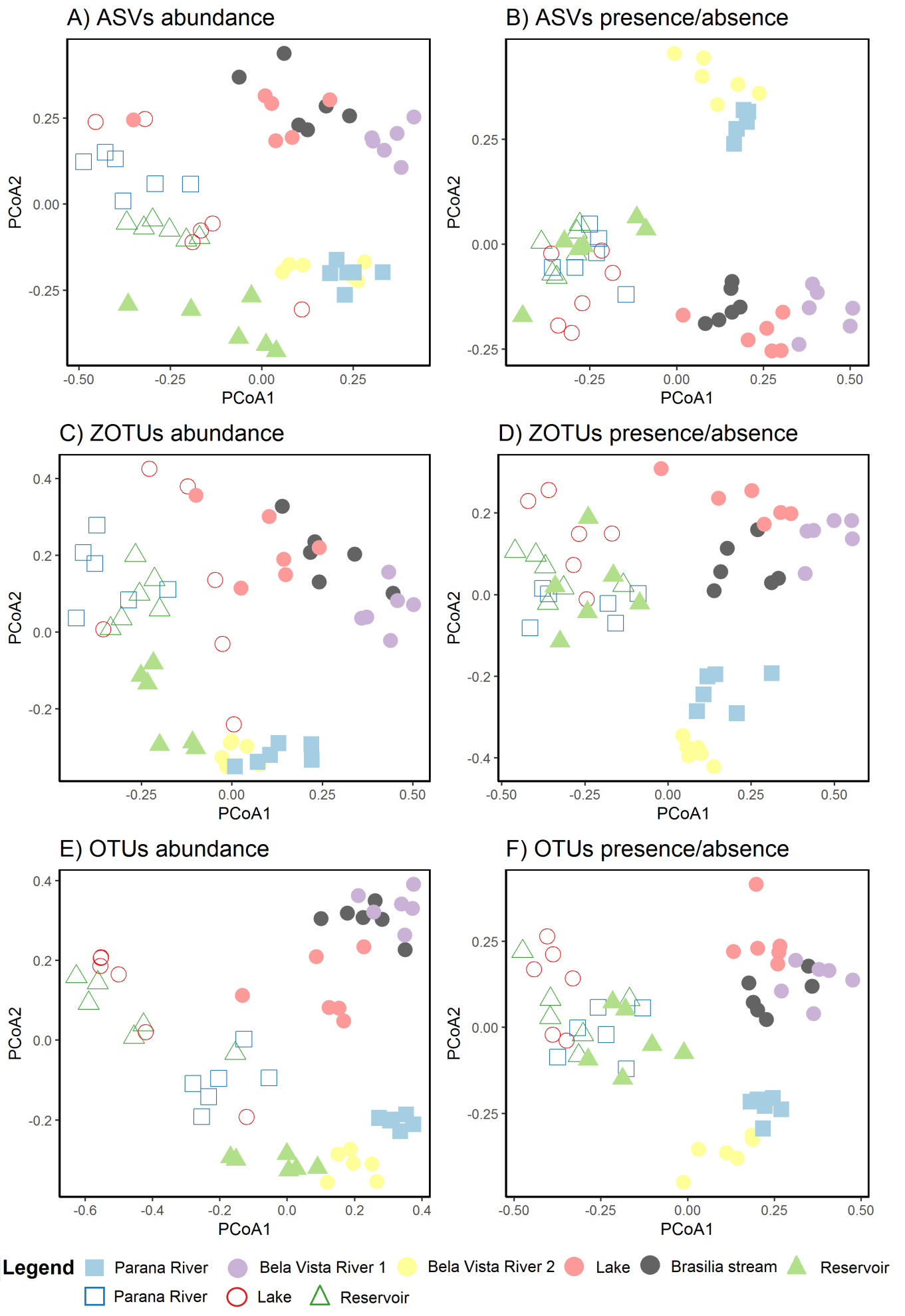

**Figure S3.** Principal Coordinates Analysis (PCoA) of fish communities from Itaipu based on sample rarefaction for A) ASVs abundance, B) ASVs presence/absence, C) ZOTUs abundance, D) ZOTUs presence/absence, E) OTUs abundance, and F) OTUs presence/absence. The axis 1 separated mainly the samples by year, while the axis 2 separated samples mainly by locality. Each point is one of the sampled replicates. Colors and symbols represent collection sites (mouth of Paraná River = blue square, Itaipu’s reservoir = green triangle, and Piracema Channel = circles [Bela Vista River 1 = purple, Bela Vista 2 = yellow, Brasilia stream = gray, and lake = red]), and tone represent year of collection (light = 2019, dark = 2020).

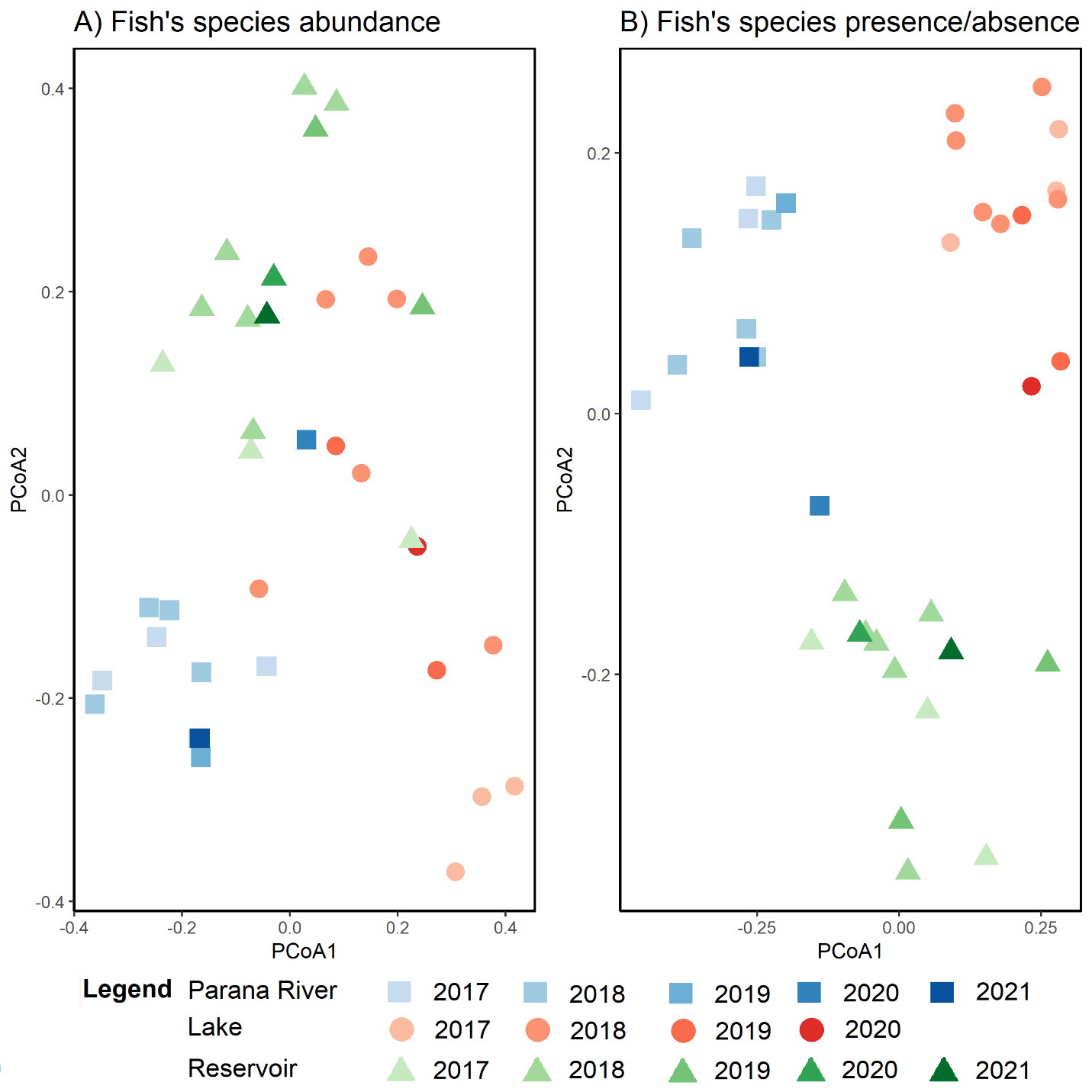

**Figure S4.** Principal Coordinates Analysis (PCoA) of fish communities from Itaipu based in traditional survey for A) abundance and B) presence/absence. The samples are clustered by locality, but not by year. Each point is one of the replicates sampled. Colors and symbols represent collection sites (mouth of Paraná River = blue square, Itaipu’s reservoir = green triangle, and lake at Piracema Channel = red circles, and tone represent year of collection (lightest = 2017 to darkest = 2021).
