## Appendix 1 for "Monitoring fish communities through environmental DNA metabarcoding in the fish pass system of the second largest hydropower plant in the world"

Itaipu metabarcoding manuscript ZOTUs Analysis


### Itaipu metabarcoding manuscript ZOTUs Analysis

###### Camila Duarte Ritter

###### July 2021

```
##   ASV_ID X10 X11 X12 X13 X14 X15 X16 X17 X18 X19 X20 X21 X22 X23 X24 X25 X26
## 1 ASV491   5 410   0   0   1   0   0   0   0   3   4   0   0   0   0   0   0
## 2 ASV588   0  13   0   0   1   0   1 196   0   2   0   0   1   1   3   0   2
## 3 ASV465   0  24   4   0   1   0   0   0   0   7   0   0   1   2 403   0   2
## 4 ASV381   6 597   2   3   4   0   1   2   1   0   2   0   1   1   2   0   2
## 5  ASV35   0   0   0   0   1   0   0   0   0   0   0   0   0   0   3   0   9
## 6  ASV84   0   0   0   0   0   1   0   2   0   0   0   0   0  15   0   4   3
##   X27 P1.1 P1.2 P1.3 P1.4 P1.5 P1.6 P2.1 P2.2 P2.3 P2.4 P2.5 P2.6 P3.1 P3.2
## 1   1    0    0    0    0    0    0    0    0    0    0    0    0    0    0
## 2   5    0    1    0    0    0    0    0    0    0    0    0    0    0    0
## 3  16    0    0    0    0    1    0    0    0    0    0    0    0    0    0
## 4   3    0    0    0    0    0    0    0    0    0    0    0    0    0    0
## 5   0    0    0    0    0    0    0    1    0    1    2    0    1    0    0
## 6  25    2    1    1    0    1    0    7    3   11    0    3    8    0    0
##   P3.3 P3.4 P3.5 P3.6 P4.1 P4.2 P4.3 P4.4 P4.5 P4.6 P5.1 P5.2 P5.3 P5.4 P5.5
## 1    0    0    0    0    0    0    0    0    0    0    0    0    0    0    0
## 2    0    0    0    0    0    0    0    0    0    0    0    0    0    0    0
## 3    0    0    0    0    0    0    0    0    0    0    0    0    0    0    0
## 4    0    0    0    0    0    0    0    0    0    0    0    0    0    0    0
## 5    0    0    0    0  307   55   66   70   73  138    0    0    0    0    1
## 6    0    0    0    0    0    4    1    3    5    5    5    6    3    1    6
##   P5.6 P6.1 P6.2 P6.3 P6.4 P6.5 P6.6
## 1    0    0    0    0    0    0    0
## 2    0    0    0    0    0    0    0
## 3    0    0    0    0    0    0    0
## 4    0    0    0    0    0    0    0
## 5    0    0    0    3    0    1    1
## 6    1   64   48   78   37   68   37
```

```
##   ZOTU_ID X10 X11 X12 X13 X14 X15 X16 X17 X18 X19 X20 X21 X22  X23 X24 X25 X26
## 1  Zotu23   0   0   1   0   0   0   0   0   0   0   0   0   0    0   1   0   7
## 2   Zotu7   0   1   2   2   0   0   0   0   1   0   0   0   6   12  20  20   4
## 3   Zotu1  27  41  22  19  15  36  20 452  11   1   3   0 119 2584  77 705 117
## 4 Zotu116   0   0   0   0   0   0   0   0   0   0   0   0   0    0   0   0   0
## 5 Zotu117   0   0   0   0   0   0   0   0   0   0   0   0   0    0   0   0   0
## 6  Zotu63   0   0   0   0   0   0   0   0   0   0   0   0   0    0   0   0   0
##    X27 P1.1 P1.2 P1.3 P1.4 P1.5 P1.6 P2.1 P2.2 P2.3 P2.4 P2.5 P2.6 P3.1 P3.2
## 1    0    0    0    0    0    0    0    1    0    1    2    0    2    0    0
## 2   10    2    0    0    0    0    0  515  874 1264  297  491 1337    1    0
## 3 1615  108  145   77    1   43   16 2347  440 1875  605  805 2730   31   28
## 4    0    0    0    0    0    0    0    0    0    0    0    0    0    0    0
## 5    0    0    0    0    0    0    0    0    0    0    0    0    0    0    0
## 6    0    0    0    0    0    2    0    0    0    0    0    0    0    0    0
##   P3.3 P3.4 P3.5 P3.6 P4.1 P4.2 P4.3 P4.4 P4.5 P4.6 P5.1 P5.2 P5.3 P5.4 P5.5
## 1    0    0    0    0  313   56   69   63   73  143    0    0    0    0    1
## 2    2    0    4    0    1    1    0    2    0    4    1    2    0    0    0
## 3    8   11   30   13 1081 1096 1328 1638 2240 1615 2246 1651  595  631 2192
## 4    0    0    0    0    0    0    0    0    0    0    0    0    0    0    0
## 5    0    0    0    0    0    0    0    0    0    0    0    0    0    0    0
## 6    0    0    0    0    2    2    1    1    2    1    0    0    0    0    0
##   P5.6  P6.1 P6.2  P6.3 P6.4  P6.5 P6.6
## 1    0     0    0     4    0     0    1
## 2    0     0    0    10    9     4    8
## 3  314 11876 8195 14749 8301 14426 8657
## 4    0   189   89   152   64    83  242
## 5    0    80   76    86   53   504   66
## 6    0   560  388   976  418   261  287
```

```
##   OUT_ID X10   X11 X12  X13   X14 X15 X16 X17 X18   X19 X20 X21  X22  X23  X24
## 1   Otu3   3    10   1    4     1   2   1   0   1     0   2   1   17   51   18
## 2   Otu8 504 54207 595 1976 12615  63  28 951 220 19326 950 396 1228 1083 4473
## 3   Otu4   3    11   2    2     0   0   0   0   1     0   0   0   10   56   20
## 4  Otu20   0     0   1    0     0   0   0   0   0     0   0   0    0    0    1
## 5   Otu1  27    41  22   19    15  36  20 452  11     1   3   0  121 2584   78
## 6  Otu97   0     0   0    0     0   0   0   0   0     0   0   0    0    0    0
##   X25  X26  X27 P1.1 P1.2 P1.3 P1.4 P1.5 P1.6 P2.1 P2.2 P2.3 P2.4 P2.5 P2.6
## 1  26   22   40    2    5    1    0    2    0 1342 2583 1803 2180 1597 3129
## 2 290 1214 3864    2    4    0    2   13    5    0    1    0    0    0    0
## 3  20    4   10    2    0    1    0    0    0  516  879 1265  299  491 1340
## 4   0    7    0    0    0    0    0    0    0    1    0    1    2    0    2
## 5 705  117 1616  108  145   77    2   45   17 2905  460 1939  655  826 2898
## 6   0    0    0    0    0    0    0    0    0    0    0    0    0    0    0
##   P3.1 P3.2 P3.3 P3.4 P3.5 P3.6 P4.1 P4.2 P4.3 P4.4 P4.5 P4.6 P5.1 P5.2 P5.3
## 1   24   41   25   11   53   20   33   43  187   45  469  110 1871 2359 1477
## 2    0    0    0    0    3    0    0    2    0    0    1    0    0    0    1
## 3    1    0    2    0    4    0    2    6    0    2    0    4    1    2    0
## 4    0    0    0    0    0    0  313   56   69   63   73  143    0    0    0
## 5   31   29    9   14   31   13 1450 1243 1487 1797 2403 1953 2278 1695  617
## 6    0    0    0    0    0    0    0    0    0    0    0    0    0    0    0
##   P5.4 P5.5 P5.6  P6.1 P6.2  P6.3 P6.4  P6.5 P6.6
## 1  729 3618  282   404  367  1019  463   651  265
## 2    1    0    0     0    0     0    0     1    0
## 3    0    0    2     1    0   114   18   103   23
## 4    0    1    0     0    0     4    0     0    1
## 5  750 2267  319 12867 8920 15659 8987 16022 9267
## 6    0    0    0    80   76    86   53   504   66
```

```
# ASVs

ASVrar <- t(ASVs_M)

Rar_ASVs <- rarecurve(ASVrar, step = 1000)
```

```
Rar_L_ASVs <- lapply(Rar_ASVs, function(x) as.data.frame(x) %>%
    tibble::rownames_to_column(var = "depth"))
names(Rar_L_ASVs) <- row.names(ASVrar)

cols <- c("#a6cee3", "#cab2d6", "#ffff99", "#fb9a99", "#636363", "#b2df8a", "#1f78b4",
    "#e31a1c", "#33a02c")

Rar_fig_ASVs <- bind_rows(Rar_L_ASVs, .id = "Sample") %>%
    mutate(depth = as.numeric(sub("N", "", depth))) %>%
    left_join(metadata) %>%
    ggplot(aes(x = depth, y = x, colour = Group, group = Sample)) + scale_color_manual(values = cols) +
    ggtitle("A) ASVs") + geom_line() + geom_vline(aes(xintercept = min(colSums(ASVs_M))),
    linetype = "dashed", colour = "darkred", size = 0.5) + labs(title = "", y = "expected ASV richness",
    x = "depth") + theme_bw() + theme(legend.position = "bottom") + theme(axis.text.x = element_text(angle = -30,
    hjust = 0)) + scale_x_continuous(labels = function(x) format(x, scientific = TRUE),
    breaks = seq(0, 1000, 1e+05))
```

```
## Joining, by = "Sample"
```

```
# ZOTUs

ZOTUrar <- t(ZOTUs_M)

Rar_ZOTUs <- rarecurve(ZOTUrar, step = 1000)
```

```
Rar_L_ZOTUs <- lapply(Rar_ZOTUs, function(x) as.data.frame(x) %>%
    tibble::rownames_to_column(var = "depth"))
names(Rar_L_ZOTUs) <- row.names(ZOTUrar)

Rar_fig_ZOTUs <- bind_rows(Rar_L_ZOTUs, .id = "Sample") %>%
    mutate(depth = as.numeric(sub("N", "", depth))) %>%
    left_join(metadata) %>%
    ggplot(aes(x = depth, y = x, colour = Group, group = Sample)) + scale_color_manual(values = cols) +
    ggtitle("B) ZOTUs") + geom_line() + geom_vline(aes(xintercept = 147), linetype = "dashed",
    colour = "darkred", size = 0.5) + labs(title = "", y = "expected ZOTU richness",
    x = "depth") + theme_bw() + theme(legend.position = "bottom") + theme(axis.text.x = element_text(angle = -30,
    hjust = 0)) + scale_x_continuous(labels = function(x) format(x, scientific = TRUE),
    breaks = seq(0, 1000, 1e+05))
```

```
## Joining, by = "Sample"
```

```
# OTUs

OTUrar <- t(OTUs_M)

Rar_OTUs <- rarecurve(OTUrar, step = 1000)
Rar_L_OTUs <- lapply(Rar_OTUs, function(x) as.data.frame(x) %>%
    tibble::rownames_to_column(var = "depth"))
names(Rar_L_OTUs) <- row.names(OTUrar)

Rar_fig_OTUs <- bind_rows(Rar_L_OTUs, .id = "Sample") %>%
    mutate(depth = as.numeric(sub("N", "", depth))) %>%
    left_join(metadata) %>%
    ggplot(aes(x = depth, y = x, colour = Group, group = Sample)) + scale_color_manual(values = cols) +
    ggtitle("C) OTUs") + geom_line() + geom_vline(aes(xintercept = min(colSums(OTUs_M))),
    linetype = "dashed", colour = "darkred", size = 0.5) + labs(title = "", y = "expected OTU richness",
    x = "depth") + theme_bw() + theme(legend.position = "bottom") + theme(axis.text.x = element_text(angle = -30,
    hjust = 0)) + scale_x_continuous(labels = function(x) format(x, scientific = TRUE),
    breaks = seq(0, 1000, 1e+05))
```

```
## Joining, by = "Sample"
```

```
# Figure combine all

newSOrder = c("Parana_River", "Bela_Vista_River2", "Bela_Vista_River1", "Lake", "Brasilia_stream",
    "Reservoir", "Parana_River2", "Lake2", "Reservoir2")

Rar_fig_ASVs$data$Group <- as.character(Rar_fig_ASVs$data$Group)
Rar_fig_ASVs$data$Group <- factor(Rar_fig_ASVs$data$Group, levels = newSOrder)

Rar_fig_ZOTUs$data$Group <- as.character(Rar_fig_ZOTUs$data$Group)
Rar_fig_ZOTUs$data$Group <- factor(Rar_fig_ZOTUs$data$Group, levels = newSOrder)

Rar_fig_OTUs$data$Group <- as.character(Rar_fig_OTUs$data$Group)
Rar_fig_OTUs$data$Group <- factor(Rar_fig_OTUs$data$Group, levels = newSOrder)

figRar <- ggarrange(Rar_fig_ASVs, Rar_fig_ZOTUs, Rar_fig_OTUs, common.legend = TRUE,
    legend = "bottom", nrow = 3)
```

```
ggsave("figRar.tiff", plot = figRar, width = 6, height = 9)
figRar
```

```
rar_Richness_ASVs <- rarefy(t(ASVs_M), sample = min(colSums(ASVs_M)), se = F)

rar_Richness_ASVs <- rar_Richness_ASVs %>%
    data.frame(Rarefied = .) %>%
    rownames_to_column(var = "Sample")

rar_Richness_ZOTUs <- rarefy(t(ZOTUs_M), sample = 147, se = F)
```

```
## Warning in rarefy(t(ZOTUs_M), sample = 147, se = F): requested 'sample' was
## larger than smallest site maximum (9)
```

```
rar_Richness_ZOTUs <- rar_Richness_ZOTUs %>%
    data.frame(Rarefied = .) %>%
    rownames_to_column(var = "Sample")

rar_Richness_OTUs <- rarefy(t(OTUs_M), sample = min(colSums(ZOTUs_M)), se = F)

rar_Richness_OTUs <- rar_Richness_OTUs %>%
    data.frame(Rarefied = .) %>%
    rownames_to_column(var = "Sample")
```

```
MCASVs <- MetaCommunity(ASVs_M)
```

```
## Warning in FUN(newX[, i], ...): Zhang-Huang sample coverage cannot be estimated
## because one probability is over 1/2. Chao estimator is returned.

## Warning in FUN(newX[, i], ...): Zhang-Huang sample coverage cannot be estimated
## because one probability is over 1/2. Chao estimator is returned.

## Warning in FUN(newX[, i], ...): Zhang-Huang sample coverage cannot be estimated
## because one probability is over 1/2. Chao estimator is returned.

## Warning in FUN(newX[, i], ...): Zhang-Huang sample coverage cannot be estimated
## because one probability is over 1/2. Chao estimator is returned.

## Warning in FUN(newX[, i], ...): Zhang-Huang sample coverage cannot be estimated
## because one probability is over 1/2. Chao estimator is returned.

## Warning in FUN(newX[, i], ...): Zhang-Huang sample coverage cannot be estimated
## because one probability is over 1/2. Chao estimator is returned.

## Warning in FUN(newX[, i], ...): Zhang-Huang sample coverage cannot be estimated
## because one probability is over 1/2. Chao estimator is returned.

## Warning in FUN(newX[, i], ...): Zhang-Huang sample coverage cannot be estimated
## because one probability is over 1/2. Chao estimator is returned.

## Warning in FUN(newX[, i], ...): Zhang-Huang sample coverage cannot be estimated
## because one probability is over 1/2. Chao estimator is returned.

## Warning in FUN(newX[, i], ...): Zhang-Huang sample coverage cannot be estimated
## because one probability is over 1/2. Chao estimator is returned.
```

```
div_ASVs <- AlphaDiversity(MCASVs, q = 1)$Communities %>%
    data.frame(effN = .) %>%
    rownames_to_column(var = "Sample")
```

```
## Warning in Coverage(Ns, Estimator = CEstimator): Zhang-Huang sample coverage
## cannot be estimated because one probability is over 1/2. Chao estimator is
## returned.
```

```
## Warning in Coverage(Ns, Estimator = CEstimator): Zhang-Huang sample coverage
## cannot be estimated because one probability is over 1/2. Chao estimator is
## returned.

## Warning in Coverage(Ns, Estimator = CEstimator): Zhang-Huang sample coverage
## cannot be estimated because one probability is over 1/2. Chao estimator is
## returned.

## Warning in Coverage(Ns, Estimator = CEstimator): Zhang-Huang sample coverage
## cannot be estimated because one probability is over 1/2. Chao estimator is
## returned.

## Warning in Coverage(Ns, Estimator = CEstimator): Zhang-Huang sample coverage
## cannot be estimated because one probability is over 1/2. Chao estimator is
## returned.

## Warning in Coverage(Ns, Estimator = CEstimator): Zhang-Huang sample coverage
## cannot be estimated because one probability is over 1/2. Chao estimator is
## returned.

## Warning in Coverage(Ns, Estimator = CEstimator): Zhang-Huang sample coverage
## cannot be estimated because one probability is over 1/2. Chao estimator is
## returned.

## Warning in Coverage(Ns, Estimator = CEstimator): Zhang-Huang sample coverage
## cannot be estimated because one probability is over 1/2. Chao estimator is
## returned.

## Warning in Coverage(Ns, Estimator = CEstimator): Zhang-Huang sample coverage
## cannot be estimated because one probability is over 1/2. Chao estimator is
## returned.

## Warning in Coverage(Ns, Estimator = CEstimator): Zhang-Huang sample coverage
## cannot be estimated because one probability is over 1/2. Chao estimator is
## returned.
```

```
MCZOTUs <- MetaCommunity(ZOTUs_M)
```

```
## Warning in FUN(newX[, i], ...): Zhang-Huang sample coverage cannot be estimated
## because one probability is over 1/2. Chao estimator is returned.
```

```
## Warning in FUN(newX[, i], ...): Zhang-Huang sample coverage cannot be estimated
## because one probability is over 1/2. Chao estimator is returned.

## Warning in FUN(newX[, i], ...): Zhang-Huang sample coverage cannot be estimated
## because one probability is over 1/2. Chao estimator is returned.

## Warning in FUN(newX[, i], ...): Zhang-Huang sample coverage cannot be estimated
## because one probability is over 1/2. Chao estimator is returned.

## Warning in FUN(newX[, i], ...): Zhang-Huang sample coverage cannot be estimated
## because one probability is over 1/2. Chao estimator is returned.

## Warning in FUN(newX[, i], ...): Zhang-Huang sample coverage cannot be estimated
## because one probability is over 1/2. Chao estimator is returned.

## Warning in FUN(newX[, i], ...): Zhang-Huang sample coverage cannot be estimated
## because one probability is over 1/2. Chao estimator is returned.

## Warning in FUN(newX[, i], ...): Zhang-Huang sample coverage cannot be estimated
## because one probability is over 1/2. Chao estimator is returned.

## Warning in FUN(newX[, i], ...): Zhang-Huang sample coverage cannot be estimated
## because one probability is over 1/2. Chao estimator is returned.

## Warning in FUN(newX[, i], ...): Zhang-Huang sample coverage cannot be estimated
## because one probability is over 1/2. Chao estimator is returned.

## Warning in FUN(newX[, i], ...): Zhang-Huang sample coverage cannot be estimated
## because one probability is over 1/2. Chao estimator is returned.

## Warning in FUN(newX[, i], ...): Zhang-Huang sample coverage cannot be estimated
## because one probability is over 1/2. Chao estimator is returned.

## Warning in FUN(newX[, i], ...): Zhang-Huang sample coverage cannot be estimated
## because one probability is over 1/2. Chao estimator is returned.
```

```
div_ZOTUs <- AlphaDiversity(MCZOTUs, q = 1)$Communities %>%
    data.frame(effN = .) %>%
    rownames_to_column(var = "Sample")
```

```
## Warning in Coverage(Ns, Estimator = CEstimator): Zhang-Huang sample coverage
## cannot be estimated because one probability is over 1/2. Chao estimator is
## returned.
```

```
## Warning in Coverage(Ns, Estimator = CEstimator): Zhang-Huang sample coverage
## cannot be estimated because one probability is over 1/2. Chao estimator is
## returned.

## Warning in Coverage(Ns, Estimator = CEstimator): Zhang-Huang sample coverage
## cannot be estimated because one probability is over 1/2. Chao estimator is
## returned.

## Warning in Coverage(Ns, Estimator = CEstimator): Zhang-Huang sample coverage
## cannot be estimated because one probability is over 1/2. Chao estimator is
## returned.

## Warning in Coverage(Ns, Estimator = CEstimator): Zhang-Huang sample coverage
## cannot be estimated because one probability is over 1/2. Chao estimator is
## returned.

## Warning in Coverage(Ns, Estimator = CEstimator): Zhang-Huang sample coverage
## cannot be estimated because one probability is over 1/2. Chao estimator is
## returned.

## Warning in Coverage(Ns, Estimator = CEstimator): Zhang-Huang sample coverage
## cannot be estimated because one probability is over 1/2. Chao estimator is
## returned.

## Warning in Coverage(Ns, Estimator = CEstimator): Zhang-Huang sample coverage
## cannot be estimated because one probability is over 1/2. Chao estimator is
## returned.

## Warning in Coverage(Ns, Estimator = CEstimator): Zhang-Huang sample coverage
## cannot be estimated because one probability is over 1/2. Chao estimator is
## returned.

## Warning in Coverage(Ns, Estimator = CEstimator): Zhang-Huang sample coverage
## cannot be estimated because one probability is over 1/2. Chao estimator is
## returned.

## Warning in Coverage(Ns, Estimator = CEstimator): Zhang-Huang sample coverage
## cannot be estimated because one probability is over 1/2. Chao estimator is
## returned.

## Warning in Coverage(Ns, Estimator = CEstimator): Zhang-Huang sample coverage
## cannot be estimated because one probability is over 1/2. Chao estimator is
## returned.

## Warning in Coverage(Ns, Estimator = CEstimator): Zhang-Huang sample coverage
## cannot be estimated because one probability is over 1/2. Chao estimator is
## returned.
```

```
MCOTUs <- MetaCommunity(OTUs_M)
```

```
## Warning in FUN(newX[, i], ...): Zhang-Huang sample coverage cannot be estimated
## because one probability is over 1/2. Chao estimator is returned.
```

```
## Warning in FUN(newX[, i], ...): Zhang-Huang sample coverage cannot be estimated
## because one probability is over 1/2. Chao estimator is returned.

## Warning in FUN(newX[, i], ...): Zhang-Huang sample coverage cannot be estimated
## because one probability is over 1/2. Chao estimator is returned.

## Warning in FUN(newX[, i], ...): Zhang-Huang sample coverage cannot be estimated
## because one probability is over 1/2. Chao estimator is returned.

## Warning in FUN(newX[, i], ...): Zhang-Huang sample coverage cannot be estimated
## because one probability is over 1/2. Chao estimator is returned.

## Warning in FUN(newX[, i], ...): Zhang-Huang sample coverage cannot be estimated
## because one probability is over 1/2. Chao estimator is returned.

## Warning in FUN(newX[, i], ...): Zhang-Huang sample coverage cannot be estimated
## because one probability is over 1/2. Chao estimator is returned.

## Warning in FUN(newX[, i], ...): Zhang-Huang sample coverage cannot be estimated
## because one probability is over 1/2. Chao estimator is returned.

## Warning in FUN(newX[, i], ...): Zhang-Huang sample coverage cannot be estimated
## because one probability is over 1/2. Chao estimator is returned.

## Warning in FUN(newX[, i], ...): Zhang-Huang sample coverage cannot be estimated
## because one probability is over 1/2. Chao estimator is returned.

## Warning in FUN(newX[, i], ...): Zhang-Huang sample coverage cannot be estimated
## because one probability is over 1/2. Chao estimator is returned.

## Warning in FUN(newX[, i], ...): Zhang-Huang sample coverage cannot be estimated
## because one probability is over 1/2. Chao estimator is returned.
```

```
div_OTUs <- AlphaDiversity(MCOTUs, q = 1)$Communities %>%
    data.frame(effN = .) %>%
    rownames_to_column(var = "Sample")
```

```
## Warning in Coverage(Ns, Estimator = CEstimator): Zhang-Huang sample coverage
## cannot be estimated because one probability is over 1/2. Chao estimator is
## returned.
```

```
## Warning in Coverage(Ns, Estimator = CEstimator): Zhang-Huang sample coverage
## cannot be estimated because one probability is over 1/2. Chao estimator is
## returned.

## Warning in Coverage(Ns, Estimator = CEstimator): Zhang-Huang sample coverage
## cannot be estimated because one probability is over 1/2. Chao estimator is
## returned.

## Warning in Coverage(Ns, Estimator = CEstimator): Zhang-Huang sample coverage
## cannot be estimated because one probability is over 1/2. Chao estimator is
## returned.

## Warning in Coverage(Ns, Estimator = CEstimator): Zhang-Huang sample coverage
## cannot be estimated because one probability is over 1/2. Chao estimator is
## returned.

## Warning in Coverage(Ns, Estimator = CEstimator): Zhang-Huang sample coverage
## cannot be estimated because one probability is over 1/2. Chao estimator is
## returned.

## Warning in Coverage(Ns, Estimator = CEstimator): Zhang-Huang sample coverage
## cannot be estimated because one probability is over 1/2. Chao estimator is
## returned.

## Warning in Coverage(Ns, Estimator = CEstimator): Zhang-Huang sample coverage
## cannot be estimated because one probability is over 1/2. Chao estimator is
## returned.

## Warning in Coverage(Ns, Estimator = CEstimator): Zhang-Huang sample coverage
## cannot be estimated because one probability is over 1/2. Chao estimator is
## returned.

## Warning in Coverage(Ns, Estimator = CEstimator): Zhang-Huang sample coverage
## cannot be estimated because one probability is over 1/2. Chao estimator is
## returned.

## Warning in Coverage(Ns, Estimator = CEstimator): Zhang-Huang sample coverage
## cannot be estimated because one probability is over 1/2. Chao estimator is
## returned.

## Warning in Coverage(Ns, Estimator = CEstimator): Zhang-Huang sample coverage
## cannot be estimated because one probability is over 1/2. Chao estimator is
## returned.
```

```
abuASVs <- phyloseq(otu_table(ASVs_M, taxa_are_rows = TRUE), sample_data(metadata))

phylodivASVs <- estimate_richness(abuASVs, split = TRUE, measures = c("Observed",
    "Chao1", "Shannon")) %>%
    rownames_to_column(var = "Sample")

abuZOTUs <- phyloseq(otu_table(ZOTUs_M, taxa_are_rows = TRUE), sample_data(metadata))

phylodivZOTUs <- estimate_richness(abuZOTUs, split = TRUE, measures = c("Observed",
    "Chao1", "Shannon")) %>%
    rownames_to_column(var = "Sample")

abuOTUs <- phyloseq(otu_table(OTUs_M, taxa_are_rows = TRUE), sample_data(metadata))

phylodivOTUs <- estimate_richness(abuOTUs, split = TRUE, measures = c("Observed",
    "Chao1", "Shannon")) %>%
    rownames_to_column(var = "Sample")
```

```
AlphaASVs <- full_join(phylodivASVs, div_ASVs, by = "Sample")
AlphaASVs <- full_join(AlphaASVs, rar_Richness_ASVs, by = "Sample")
AlphaASVs <- full_join(metadata, AlphaASVs, by = "Sample")
write.csv(AlphaASVs, "AlphaASVs.csv")

AlphaZOTUs <- full_join(phylodivZOTUs, div_ZOTUs, by = "Sample")
AlphaZOTUs <- full_join(AlphaZOTUs, rar_Richness_ZOTUs, by = "Sample")
AlphaZOTUs <- full_join(metadata, AlphaZOTUs, by = "Sample")
write.csv(AlphaZOTUs, "AlphaZOTUs.csv")

AlphaOTUs <- full_join(phylodivOTUs, div_OTUs, by = "Sample")
AlphaOTUs <- full_join(AlphaOTUs, rar_Richness_OTUs, by = "Sample")
AlphaOTUs <- full_join(metadata, AlphaOTUs, by = "Sample")
write.csv(AlphaOTUs, "AlphaOTUs.csv")
```

```
forms <- c(15, 16, 16, 16, 16, 17, 0, 1, 2)

newSOrder2 = c("Parana_River_mouth", "Bela_Vista_River2", "Bela_Vista_River1", "Lake",
    "Brasilia_stream", "Reservoir", "Parana_River_mouth2", "Lake2", "Reservoir2")

AlphaASVs2 <- read.csv("AlphaASVs2.csv")

AlphaASVsplot <- ggplot(AlphaASVs2) + aes(x = Group, y = Alpha, colour = Group, shape = Group) +
    geom_point(size = 3) + scale_color_manual(values = cols) + scale_shape_manual(values = forms) +
    ggtitle("A) ASVs") + theme(axis.text.x = element_blank(), axis.ticks.x = element_blank(),
    plot.title = element_text(size = 15L), plot.caption = element_text(size = 12L),
    panel.grid.major.x = element_line(size = 0.5, colour = "#f0f0f0"), panel.grid.major.y = element_blank(),
    panel.grid.minor = element_blank(), panel.background = element_blank(), axis.line = element_line(colour = "black"),
    panel.border = element_rect(colour = "black", fill = NA, size = 1), legend.position = ("botton")) +
    facet_wrap(vars(Measure), ncol = 5, scales = "free")

AlphaZOTUs2 <- read.csv("AlphaZOTUs2.csv")

AlphaZOTUsplot <- ggplot(AlphaZOTUs2) + aes(x = Group, y = Alpha, colour = Group,
    shape = Group) + geom_point(size = 3) + scale_color_manual(values = cols) + scale_shape_manual(values = forms) +
    ggtitle("B) ZOTUs") + theme(axis.text.x = element_blank(), axis.ticks.x = element_blank(),
    plot.title = element_text(size = 15L), plot.caption = element_text(size = 12L),
    panel.grid.major.x = element_line(size = 0.5, colour = "#f0f0f0"), panel.grid.major.y = element_blank(),
    panel.grid.minor = element_blank(), panel.background = element_blank(), axis.line = element_line(colour = "black"),
    panel.border = element_rect(colour = "black", fill = NA, size = 1), legend.position = ("botton")) +
    facet_wrap(vars(Measure), ncol = 5, scales = "free")

AlphaOTUs2 <- read.csv("AlphaOTUs2.csv")

AlphaOTUsplot <- ggplot(AlphaOTUs2) + aes(x = Group, y = Alpha, colour = Group, shape = Group) +
    geom_point(size = 3) + scale_color_manual(values = cols) + scale_shape_manual(values = forms) +
    ggtitle("C) OTUs") + theme(axis.text.x = element_blank(), axis.ticks.x = element_blank(),
    plot.title = element_text(size = 15L), plot.caption = element_text(size = 12L),
    panel.grid.major.x = element_line(size = 0.5, colour = "#f0f0f0"), panel.grid.major.y = element_blank(),
    panel.grid.minor = element_blank(), panel.background = element_blank(), axis.line = element_line(colour = "black"),
    panel.border = element_rect(colour = "black", fill = NA, size = 1), legend.position = ("botton")) +
    facet_wrap(vars(Measure), ncol = 5, scales = "free")

AlphaASVsplot$data$Group <- as.character(AlphaASVsplot$data$Group)
AlphaASVsplot$data$Group <- factor(AlphaASVsplot$data$Group, levels = newSOrder2)

AlphaZOTUsplot$data$Group <- as.character(AlphaZOTUsplot$data$Group)
AlphaZOTUsplot$data$Group <- factor(AlphaZOTUsplot$data$Group, levels = newSOrder2)

AlphaOTUsplot$data$Group <- as.character(AlphaOTUsplot$data$Group)
AlphaOTUsplot$data$Group <- factor(AlphaOTUsplot$data$Group, levels = newSOrder2)

figAlpha <- ggarrange(AlphaASVsplot, AlphaZOTUsplot, AlphaOTUsplot, common.legend = TRUE,
    legend = "bottom", nrow = 3)
ggsave("figAlpha.tiff", figAlpha, width = 8, height = 12, device = tiff)
figAlpha
```

```
# PCoA abu

PcoA_ASVs <- cmdscale(distASVs, k = (nrow(metadata) - 1), eig = TRUE)
```

```
## Warning in cmdscale(distASVs, k = (nrow(metadata) - 1), eig = TRUE): only 41 of
## the first 53 eigenvalues are > 0
```

```
PC_ASVs <- data.frame(scores(PcoA_ASVs))
PC_ASVs$Sample <- rownames(PC_ASVs)

PC_ASVs <- PC_ASVs %>%
    as_data_frame() %>%
    inner_join(metadata, by = "Sample")
```

```
## Warning: `as_data_frame()` was deprecated in tibble 2.0.0.
## Please use `as_tibble()` instead.
## The signature and semantics have changed, see `?as_tibble`.
## This warning is displayed once every 8 hours.
## Call `lifecycle::last_lifecycle_warnings()` to see where this warning was generated.
```

```
# head(PC_ASVs)

## envfit

envfit(PC_ASVs[, 1:2] ~ as.factor(PC_ASVs$Local))
```

```
## 
## ***FACTORS:
## 
## Centroids:
##                                              Dim1    Dim2
## as.factor(PC_ASVs$Local)Bela_Vista_River1  0.1893  0.3363
## as.factor(PC_ASVs$Local)Bela_Vista_River2  0.3104 -0.2792
## as.factor(PC_ASVs$Local)Brasilia_stream    0.2272 -0.0634
## as.factor(PC_ASVs$Local)Lake              -0.1408 -0.1793
## as.factor(PC_ASVs$Local)Parana_River       0.2421  0.2128
## as.factor(PC_ASVs$Local)Parana_River2     -0.0943  0.0898
## as.factor(PC_ASVs$Local)Reservoir         -0.2966  0.0312
## 
## Goodness of fit:
##                              r2 Pr(>r)    
## as.factor(PC_ASVs$Local) 0.8283  0.001 ***
## ---
## Signif. codes:  0 '***' 0.001 '**' 0.01 '*' 0.05 '.' 0.1 ' ' 1
## Permutation: free
## Number of permutations: 999
```

```
envfit(PC_ASVs[, 1:2] ~ as.factor(PC_ASVs$Year))
```

```
## 
## ***FACTORS:
## 
## Centroids:
##                                    Dim1    Dim2
## as.factor(PC_ASVs$Year)Nineteen  0.1339 -0.0096
## as.factor(PC_ASVs$Year)Twenty   -0.2677  0.0191
## 
## Goodness of fit:
##                             r2 Pr(>r)    
## as.factor(PC_ASVs$Year) 0.3435  0.001 ***
## ---
## Signif. codes:  0 '***' 0.001 '**' 0.01 '*' 0.05 '.' 0.1 ' ' 1
## Permutation: free
## Number of permutations: 999
```

```
# PCoA presence/absence

PcoA_ASVspa <- cmdscale(distASVspa, k = (nrow(metadata) - 1), eig = TRUE)
```

```
## Warning in cmdscale(distASVspa, k = (nrow(metadata) - 1), eig = TRUE): only 31
## of the first 53 eigenvalues are > 0
```

```
PC_ASVspa <- data.frame(scores(PcoA_ASVspa))
PC_ASVspa$Sample <- rownames(PC_ASVspa)

PC_ASVspa <- PC_ASVspa %>%
    as_data_frame() %>%
    inner_join(metadata, by = "Sample")
# head(PC_ASVspa)

## envfit

envfit(PC_ASVspa[, 1:2] ~ as.factor(PC_ASVspa$Local))
```

```
## 
## ***FACTORS:
## 
## Centroids:
##                                                Dim1    Dim2
## as.factor(PC_ASVspa$Local)Bela_Vista_River1  0.2228  0.2195
## as.factor(PC_ASVspa$Local)Bela_Vista_River2  0.2239 -0.2344
## as.factor(PC_ASVspa$Local)Brasilia_stream    0.2033  0.0503
## as.factor(PC_ASVspa$Local)Lake              -0.1706 -0.1745
## as.factor(PC_ASVspa$Local)Parana_River       0.2549  0.1253
## as.factor(PC_ASVspa$Local)Parana_River2     -0.0598  0.1292
## as.factor(PC_ASVspa$Local)Reservoir         -0.2520  0.0296
## 
## Goodness of fit:
##                                r2 Pr(>r)    
## as.factor(PC_ASVspa$Local) 0.7083  0.001 ***
## ---
## Signif. codes:  0 '***' 0.001 '**' 0.01 '*' 0.05 '.' 0.1 ' ' 1
## Permutation: free
## Number of permutations: 999
```

```
envfit(PC_ASVspa[, 1:2] ~ as.factor(PC_ASVspa$Year))
```

```
## 
## ***FACTORS:
## 
## Centroids:
##                                      Dim1    Dim2
## as.factor(PC_ASVspa$Year)Nineteen  0.1354 -0.0305
## as.factor(PC_ASVspa$Year)Twenty   -0.2709  0.0610
## 
## Goodness of fit:
##                               r2 Pr(>r)    
## as.factor(PC_ASVspa$Year) 0.4125  0.001 ***
## ---
## Signif. codes:  0 '***' 0.001 '**' 0.01 '*' 0.05 '.' 0.1 ' ' 1
## Permutation: free
## Number of permutations: 999
```

```
# PCoA abu

PcoA_ZOTUs <- cmdscale(distZOTUs, k = (50), eig = TRUE)
```

```
## Warning in cmdscale(distZOTUs, k = (50), eig = TRUE): only 35 of the first 50
## eigenvalues are > 0
```

```
PC_ZOTUs <- data.frame(scores(PcoA_ZOTUs))
PC_ZOTUs$Sample <- rownames(PC_ZOTUs)

PC_ZOTUs <- PC_ZOTUs %>%
    as_data_frame() %>%
    inner_join(metadata, by = "Sample")
# head(PC_ZOTUs)

## envfit

envfit(PC_ZOTUs[, 1:2] ~ as.factor(PC_ZOTUs$Local))
```

```
## 
## ***FACTORS:
## 
## Centroids:
##                                               Dim1    Dim2
## as.factor(PC_ZOTUs$Local)Bela_Vista_River1  0.0878  0.3712
## as.factor(PC_ZOTUs$Local)Bela_Vista_River2  0.3822 -0.2161
## as.factor(PC_ZOTUs$Local)Brasilia_stream    0.2430 -0.0190
## as.factor(PC_ZOTUs$Local)Lake              -0.0782 -0.1545
## as.factor(PC_ZOTUs$Local)Parana_River       0.1468  0.2515
## as.factor(PC_ZOTUs$Local)Parana_River2     -0.0945 -0.0147
## as.factor(PC_ZOTUs$Local)Reservoir         -0.3045 -0.0320
## 
## Goodness of fit:
##                               r2 Pr(>r)    
## as.factor(PC_ZOTUs$Local) 0.8257  0.001 ***
## ---
## Signif. codes:  0 '***' 0.001 '**' 0.01 '*' 0.05 '.' 0.1 ' ' 1
## Permutation: free
## Number of permutations: 999
```

```
envfit(PC_ZOTUs[, 1:2] ~ as.factor(PC_ZOTUs$Year))
```

```
## 
## ***FACTORS:
## 
## Centroids:
##                                     Dim1    Dim2
## as.factor(PC_ZOTUs$Year)Nineteen  0.1208  0.0218
## as.factor(PC_ZOTUs$Year)Twenty   -0.2416 -0.0437
## 
## Goodness of fit:
##                              r2 Pr(>r)    
## as.factor(PC_ZOTUs$Year) 0.3031  0.001 ***
## ---
## Signif. codes:  0 '***' 0.001 '**' 0.01 '*' 0.05 '.' 0.1 ' ' 1
## Permutation: free
## Number of permutations: 999
```

```
# PCoA presence/absence

PcoA_ZOTUspa <- cmdscale(distZOTUspa, k = (50), eig = TRUE)
```

```
## Warning in cmdscale(distZOTUspa, k = (50), eig = TRUE): only 28 of the first 50
## eigenvalues are > 0
```

```
PC_ZOTUspa <- data.frame(scores(PcoA_ZOTUspa))
PC_ZOTUspa$Sample <- rownames(PC_ZOTUspa)

PC_ZOTUspa <- PC_ZOTUspa %>%
    as_data_frame() %>%
    inner_join(metadata, by = "Sample")
# head(PC_ZOTUspa)

## envfit

envfit(PC_ZOTUspa[, 1:2] ~ as.factor(PC_ZOTUspa$Local))
```

```
## 
## ***FACTORS:
## 
## Centroids:
##                                                 Dim1    Dim2
## as.factor(PC_ZOTUspa$Local)Bela_Vista_River1  0.2211  0.2643
## as.factor(PC_ZOTUspa$Local)Bela_Vista_River2  0.2452 -0.1969
## as.factor(PC_ZOTUspa$Local)Brasilia_stream    0.1937  0.0020
## as.factor(PC_ZOTUspa$Local)Lake              -0.1465 -0.1003
## as.factor(PC_ZOTUspa$Local)Parana_River       0.2056  0.1559
## as.factor(PC_ZOTUspa$Local)Parana_River2     -0.0192 -0.0583
## as.factor(PC_ZOTUspa$Local)Reservoir         -0.2767  0.0168
## 
## Goodness of fit:
##                                 r2 Pr(>r)    
## as.factor(PC_ZOTUspa$Local) 0.7178  0.001 ***
## ---
## Signif. codes:  0 '***' 0.001 '**' 0.01 '*' 0.05 '.' 0.1 ' ' 1
## Permutation: free
## Number of permutations: 999
```

```
envfit(PC_ZOTUspa[, 1:2] ~ as.factor(PC_ZOTUspa$Year))
```

```
## 
## ***FACTORS:
## 
## Centroids:
##                                       Dim1    Dim2
## as.factor(PC_ZOTUspa$Year)Nineteen  0.1123 -0.0062
## as.factor(PC_ZOTUspa$Year)Twenty   -0.2246  0.0123
## 
## Goodness of fit:
##                                r2 Pr(>r)    
## as.factor(PC_ZOTUspa$Year) 0.3014  0.001 ***
## ---
## Signif. codes:  0 '***' 0.001 '**' 0.01 '*' 0.05 '.' 0.1 ' ' 1
## Permutation: free
## Number of permutations: 999
```

```
# PCoA abu

PcoA_OTUs <- cmdscale(distOTUs, k = (50), eig = TRUE)
```

```
## Warning in cmdscale(distOTUs, k = (50), eig = TRUE): only 35 of the first 50
## eigenvalues are > 0
```

```
PC_OTUs <- data.frame(scores(PcoA_OTUs))
PC_OTUs$Sample <- rownames(PC_OTUs)

PC_OTUs <- PC_OTUs %>%
    as_data_frame() %>%
    inner_join(metadata, by = "Sample")
# head(PC_ZOTUs)

## envfit

envfit(PC_OTUs[, 1:2] ~ as.factor(PC_OTUs$Local))
```

```
## 
## ***FACTORS:
## 
## Centroids:
##                                              Dim1    Dim2
## as.factor(PC_OTUs$Local)Bela_Vista_River1  0.1280 -0.3321
## as.factor(PC_OTUs$Local)Bela_Vista_River2  0.3510  0.2520
## as.factor(PC_OTUs$Local)Brasilia_stream    0.2637 -0.0225
## as.factor(PC_OTUs$Local)Lake              -0.1129  0.1585
## as.factor(PC_OTUs$Local)Parana_River       0.1784 -0.2190
## as.factor(PC_OTUs$Local)Parana_River2     -0.1261 -0.0238
## as.factor(PC_OTUs$Local)Reservoir         -0.2846  0.0143
## 
## Goodness of fit:
##                              r2 Pr(>r)    
## as.factor(PC_OTUs$Local) 0.8207  0.001 ***
## ---
## Signif. codes:  0 '***' 0.001 '**' 0.01 '*' 0.05 '.' 0.1 ' ' 1
## Permutation: free
## Number of permutations: 999
```

```
envfit(PC_OTUs[, 1:2] ~ as.factor(PC_OTUs$Year))
```

```
## 
## ***FACTORS:
## 
## Centroids:
##                                    Dim1    Dim2
## as.factor(PC_OTUs$Year)Nineteen  0.1316 -0.0103
## as.factor(PC_OTUs$Year)Twenty   -0.2632  0.0206
## 
## Goodness of fit:
##                             r2 Pr(>r)    
## as.factor(PC_OTUs$Year) 0.3586  0.001 ***
## ---
## Signif. codes:  0 '***' 0.001 '**' 0.01 '*' 0.05 '.' 0.1 ' ' 1
## Permutation: free
## Number of permutations: 999
```

```
# PCoA presence/absence

PcoA_OTUspa <- cmdscale(distOTUspa, k = (50), eig = TRUE)
```

```
## Warning in cmdscale(distOTUspa, k = (50), eig = TRUE): only 28 of the first 50
## eigenvalues are > 0
```

```
PC_OTUspa <- data.frame(scores(PcoA_OTUspa))
PC_OTUspa$Sample <- rownames(PC_OTUspa)

PC_OTUspa <- PC_OTUspa %>%
    as_data_frame() %>%
    inner_join(metadata, by = "Sample")
# head(PC_OTUspa)

## envfit

envfit(PC_OTUspa[, 1:2] ~ as.factor(PC_OTUspa$Local))
```

```
## 
## ***FACTORS:
## 
## Centroids:
##                                                Dim1    Dim2
## as.factor(PC_OTUspa$Local)Bela_Vista_River1  0.2056  0.2202
## as.factor(PC_OTUspa$Local)Bela_Vista_River2  0.2378 -0.2345
## as.factor(PC_OTUspa$Local)Brasilia_stream    0.2129  0.0542
## as.factor(PC_OTUspa$Local)Lake              -0.1538 -0.0960
## as.factor(PC_OTUspa$Local)Parana_River       0.1962  0.1563
## as.factor(PC_OTUspa$Local)Parana_River2     -0.0587 -0.0252
## as.factor(PC_OTUspa$Local)Reservoir         -0.2431  0.0105
## 
## Goodness of fit:
##                                r2 Pr(>r)    
## as.factor(PC_OTUspa$Local) 0.7298  0.001 ***
## ---
## Signif. codes:  0 '***' 0.001 '**' 0.01 '*' 0.05 '.' 0.1 ' ' 1
## Permutation: free
## Number of permutations: 999
```

```
envfit(PC_OTUspa[, 1:2] ~ as.factor(PC_OTUspa$Year))
```

```
## 
## ***FACTORS:
## 
## Centroids:
##                                      Dim1    Dim2
## as.factor(PC_OTUspa$Year)Nineteen  0.1101 -0.0098
## as.factor(PC_OTUspa$Year)Twenty   -0.2202  0.0196
## 
## Goodness of fit:
##                             r2 Pr(>r)    
## as.factor(PC_OTUspa$Year) 0.32  0.001 ***
## ---
## Signif. codes:  0 '***' 0.001 '**' 0.01 '*' 0.05 '.' 0.1 ' ' 1
## Permutation: free
## Number of permutations: 999
```

```
# Abus hill number

figPCoAASVs <- ggplot(PC_ASVs) + aes(x = Dim1, y = Dim2, color = Group, shape = Group,
    size = 3) + xlab("PCoA1") + ylab("PCoA2") + scale_fill_manual(values = cols) +
    scale_color_manual(values = cols) + scale_shape_manual(values = forms) + ggtitle("A) ASVs abundance") +
    geom_point(position = position_jitter(0.1)) + theme(panel.grid.major = element_blank(),
    panel.grid.minor = element_blank(), panel.background = element_blank(), plot.title = element_text(size = 15L),
    plot.caption = element_text(size = 12L), axis.line = element_line(colour = "black"),
    panel.border = element_rect(colour = "black", fill = NA, size = 1))

figPCoAZOTUs <- ggplot(PC_ZOTUs) + aes(x = Dim1, y = Dim2, color = Group, shape = Group,
    size = 3) + xlab("PCoA1") + ylab("PCoA2") + scale_fill_manual(values = cols) +
    scale_color_manual(values = cols) + scale_shape_manual(values = forms) + ggtitle("C) ZOTUs abundance") +
    geom_point(position = position_jitter(0.1)) + theme(panel.grid.major = element_blank(),
    panel.grid.minor = element_blank(), panel.background = element_blank(), plot.title = element_text(size = 15L),
    plot.caption = element_text(size = 12L), axis.line = element_line(colour = "black"),
    panel.border = element_rect(colour = "black", fill = NA, size = 1))

figPCoAOTUs <- ggplot(PC_OTUs) + aes(x = Dim1, y = Dim2, color = Group, shape = Group,
    size = 3) + xlab("PCoA1") + ylab("PCoA2") + scale_fill_manual(values = cols) +
    scale_color_manual(values = cols) + scale_shape_manual(values = forms) + ggtitle("E) OTUs abundance") +
    geom_point(position = position_jitter(0.1)) + theme(panel.grid.major = element_blank(),
    panel.grid.minor = element_blank(), panel.background = element_blank(), plot.title = element_text(size = 15L),
    plot.caption = element_text(size = 12L), axis.line = element_line(colour = "black"),
    panel.border = element_rect(colour = "black", fill = NA, size = 1))

# presence/absence hill number

figPCoAASVspa <- ggplot(PC_ASVspa) + aes(x = Dim1, y = Dim2, color = Group, shape = Group,
    size = 3) + xlab("PCoA1") + ylab("PCoA2") + scale_fill_manual(values = cols) +
    scale_color_manual(values = cols) + scale_shape_manual(values = forms) + ggtitle("B) ASVs presence/absence") +
    geom_point(position = position_jitter(0.1)) + theme(panel.grid.major = element_blank(),
    panel.grid.minor = element_blank(), panel.background = element_blank(), plot.title = element_text(size = 15L),
    plot.caption = element_text(size = 12L), axis.line = element_line(colour = "black"),
    panel.border = element_rect(colour = "black", fill = NA, size = 1))

figPCoAZOTUspa <- ggplot(PC_ZOTUspa) + aes(x = Dim1, y = Dim2, color = Group, shape = Group,
    size = 3) + xlab("PCoA1") + ylab("PCoA2") + scale_fill_manual(values = cols) +
    scale_color_manual(values = cols) + scale_shape_manual(values = forms) + ggtitle("D) ZOTUs presence/absence") +
    geom_point(position = position_jitter(0.1)) + theme(panel.grid.major = element_blank(),
    panel.grid.minor = element_blank(), panel.background = element_blank(), plot.title = element_text(size = 15L),
    plot.caption = element_text(size = 12L), axis.line = element_line(colour = "black"),
    panel.border = element_rect(colour = "black", fill = NA, size = 1))

figPCoAOTUspa <- ggplot(PC_OTUspa) + aes(x = Dim1, y = Dim2, color = Group, shape = Group,
    size = 3) + xlab("PCoA1") + ylab("PCoA2") + scale_fill_manual(values = cols) +
    scale_color_manual(values = cols) + scale_shape_manual(values = forms) + ggtitle("F) OTUs presence/absence") +
    geom_point(position = position_jitter(0.1)) + theme(panel.grid.major = element_blank(),
    panel.grid.minor = element_blank(), panel.background = element_blank(), plot.title = element_text(size = 15L),
    plot.caption = element_text(size = 12L), axis.line = element_line(colour = "black"),
    panel.border = element_rect(colour = "black", fill = NA, size = 1))

# Plot all

figPCoAASVs$data$Group <- as.character(figPCoAASVs$data$Group)
figPCoAASVs$data$Group <- factor(figPCoAASVs$data$Group, levels = newSOrder)

figPCoAZOTUs$data$Group <- as.character(figPCoAZOTUs$data$Group)
figPCoAZOTUs$data$Group <- factor(figPCoAZOTUs$data$Group, levels = newSOrder)

figPCoAOTUs$data$Group <- as.character(figPCoAOTUs$data$Group)
figPCoAOTUs$data$Group <- factor(figPCoAOTUs$data$Group, levels = newSOrder)

figPCoAASVspa$data$Group <- as.character(figPCoAASVspa$data$Group)
figPCoAASVspa$data$Group <- factor(figPCoAASVspa$data$Group, levels = newSOrder)

figPCoAZOTUspa$data$Group <- as.character(figPCoAZOTUspa$data$Group)
figPCoAZOTUspa$data$Group <- factor(figPCoAZOTUspa$data$Group, levels = newSOrder)

figPCoAOTUspa$data$Group <- as.character(figPCoAOTUspa$data$Group)
figPCoAOTUspa$data$Group <- factor(figPCoAOTUspa$data$Group, levels = newSOrder)


figPCoA <- ggarrange(figPCoAASVs, figPCoAASVspa, figPCoAZOTUs, figPCoAZOTUspa, figPCoAOTUs,
    figPCoAOTUspa, common.legend = TRUE, legend = "bottom", nrow = 3, ncol = 2)
ggsave("figPCoA.tiff", figPCoA, width = 8, height = 12, device = tiff)
figPCoA
```

```
# rarefy down to lowest sample
comRarASVs <- rrarefy(t(ASVs_M), sample = min(colSums(ASVs_M)))

distRarASVs <- vegdist(comRarASVs, method = "bray")
distRarASVspa <- vegdist(comRarASVs, method = "bray", binary = TRUE)

# PCoA abu

PcoA_RarASVs <- cmdscale(distRarASVs, k = (nrow(metadata) - 1), eig = TRUE)
```

```
## Warning in cmdscale(distRarASVs, k = (nrow(metadata) - 1), eig = TRUE): only 41
## of the first 53 eigenvalues are > 0
```

```
PC_RarASVs <- data.frame(scores(PcoA_RarASVs))
PC_RarASVs$Sample <- rownames(PC_RarASVs)

PC_RarASVs <- PC_RarASVs %>%
    as_data_frame() %>%
    inner_join(metadata, by = "Sample")
# head(PC_RarASVs)

## envfit

envfit(PC_RarASVs[, 1:2] ~ as.factor(PC_RarASVs$Local))
```

```
## 
## ***FACTORS:
## 
## Centroids:
##                                                 Dim1    Dim2
## as.factor(PC_RarASVs$Local)Bela_Vista_River1  0.0760 -0.3113
## as.factor(PC_RarASVs$Local)Bela_Vista_River2  0.3970  0.1851
## as.factor(PC_RarASVs$Local)Brasilia_stream    0.2023  0.2660
## as.factor(PC_RarASVs$Local)Lake              -0.0778  0.1427
## as.factor(PC_RarASVs$Local)Parana_River       0.2210 -0.2988
## as.factor(PC_RarASVs$Local)Parana_River2     -0.3199  0.0924
## as.factor(PC_RarASVs$Local)Reservoir         -0.2104 -0.1094
## 
## Goodness of fit:
##                                 r2 Pr(>r)    
## as.factor(PC_RarASVs$Local) 0.8161  0.001 ***
## ---
## Signif. codes:  0 '***' 0.001 '**' 0.01 '*' 0.05 '.' 0.1 ' ' 1
## Permutation: free
## Number of permutations: 999
```

```
envfit(PC_RarASVs[, 1:2] ~ as.factor(PC_RarASVs$Year))
```

```
## 
## ***FACTORS:
## 
## Centroids:
##                                       Dim1    Dim2
## as.factor(PC_RarASVs$Year)Nineteen  0.1287 -0.0276
## as.factor(PC_RarASVs$Year)Twenty   -0.2574  0.0553
## 
## Goodness of fit:
##                                r2 Pr(>r)    
## as.factor(PC_RarASVs$Year) 0.3102  0.001 ***
## ---
## Signif. codes:  0 '***' 0.001 '**' 0.01 '*' 0.05 '.' 0.1 ' ' 1
## Permutation: free
## Number of permutations: 999
```

```
# PCoA presence/absence

PcoA_RarASVspa <- cmdscale(distRarASVspa, k = (nrow(metadata) - 1), eig = TRUE)
```

```
## Warning in cmdscale(distRarASVspa, k = (nrow(metadata) - 1), eig = TRUE): only
## 33 of the first 53 eigenvalues are > 0
```

```
PC_RarASVspa <- data.frame(scores(PcoA_RarASVspa))
PC_RarASVspa$Sample <- rownames(PC_RarASVspa)

PC_RarASVspa <- PC_RarASVspa %>%
    as_data_frame() %>%
    inner_join(metadata, by = "Sample")
# head(PC_RarASVspa)

## envfit

envfit(PC_RarASVspa[, 1:2] ~ as.factor(PC_RarASVspa$Local))
```

```
## 
## ***FACTORS:
## 
## Centroids:
##                                                   Dim1    Dim2
## as.factor(PC_RarASVspa$Local)Bela_Vista_River1 -0.1215  0.3815
## as.factor(PC_RarASVspa$Local)Bela_Vista_River2 -0.3824 -0.2597
## as.factor(PC_RarASVspa$Local)Brasilia_stream   -0.2491 -0.0692
## as.factor(PC_RarASVspa$Local)Lake               0.0641 -0.1691
## as.factor(PC_RarASVspa$Local)Parana_River      -0.2345  0.2615
## as.factor(PC_RarASVspa$Local)Parana_River2      0.2546  0.0600
## as.factor(PC_RarASVspa$Local)Reservoir          0.3024 -0.0179
## 
## Goodness of fit:
##                                   r2 Pr(>r)    
## as.factor(PC_RarASVspa$Local) 0.8137  0.001 ***
## ---
## Signif. codes:  0 '***' 0.001 '**' 0.01 '*' 0.05 '.' 0.1 ' ' 1
## Permutation: free
## Number of permutations: 999
```

```
envfit(PC_RarASVspa[, 1:2] ~ as.factor(PC_RarASVspa$Year))
```

```
## 
## ***FACTORS:
## 
## Centroids:
##                                         Dim1    Dim2
## as.factor(PC_RarASVspa$Year)Nineteen -0.1560  0.0152
## as.factor(PC_RarASVspa$Year)Twenty    0.3121 -0.0305
## 
## Goodness of fit:
##                                  r2 Pr(>r)    
## as.factor(PC_RarASVspa$Year) 0.4084  0.001 ***
## ---
## Signif. codes:  0 '***' 0.001 '**' 0.01 '*' 0.05 '.' 0.1 ' ' 1
## Permutation: free
## Number of permutations: 999
```

```
# rarefy down to lowest sample
comRarZOTUs <- rrarefy(t(ZOTUs_M), sample = 147)
```

```
## Warning in rrarefy(t(ZOTUs_M), sample = 147): some row sums < 'sample' and are
## not rarefied
```

```
distRarZOTUs <- vegdist(comRarZOTUs, method = "bray")
distRarZOTUspa <- vegdist(comRarZOTUs, method = "bray", binary = TRUE)

# PCoA abu

PcoA_RarZOTUs <- cmdscale(distRarZOTUs, k = (50), eig = TRUE)
```

```
## Warning in cmdscale(distRarZOTUs, k = (50), eig = TRUE): only 37 of the first 50
## eigenvalues are > 0
```

```
PC_RarZOTUs <- data.frame(scores(PcoA_RarZOTUs))
PC_RarZOTUs$Sample <- rownames(PC_RarZOTUs)

PC_RarZOTUs <- PC_RarZOTUs %>%
    as_data_frame() %>%
    inner_join(metadata, by = "Sample")
# head(PC_RarZOTUs)

## envfit

envfit(PC_RarZOTUs[, 1:2] ~ as.factor(PC_RarZOTUs$Local))
```

```
## 
## ***FACTORS:
## 
## Centroids:
##                                                  Dim1    Dim2
## as.factor(PC_RarZOTUs$Local)Bela_Vista_River1 -0.0713  0.3285
## as.factor(PC_RarZOTUs$Local)Bela_Vista_River2  0.4747 -0.0510
## as.factor(PC_RarZOTUs$Local)Brasilia_stream    0.2894 -0.1939
## as.factor(PC_RarZOTUs$Local)Lake              -0.0197 -0.1524
## as.factor(PC_RarZOTUs$Local)Parana_River       0.1166  0.3383
## as.factor(PC_RarZOTUs$Local)Parana_River2     -0.3408 -0.1650
## as.factor(PC_RarZOTUs$Local)Reservoir         -0.2146  0.0240
## 
## Goodness of fit:
##                                  r2 Pr(>r)    
## as.factor(PC_RarZOTUs$Local) 0.8244  0.001 ***
## ---
## Signif. codes:  0 '***' 0.001 '**' 0.01 '*' 0.05 '.' 0.1 ' ' 1
## Permutation: free
## Number of permutations: 999
```

```
envfit(PC_RarZOTUs[, 1:2] ~ as.factor(PC_RarZOTUs$Year))
```

```
## 
## ***FACTORS:
## 
## Centroids:
##                                        Dim1    Dim2
## as.factor(PC_RarZOTUs$Year)Nineteen  0.1096  0.0710
## as.factor(PC_RarZOTUs$Year)Twenty   -0.2193 -0.1421
## 
## Goodness of fit:
##                                 r2 Pr(>r)    
## as.factor(PC_RarZOTUs$Year) 0.2897  0.001 ***
## ---
## Signif. codes:  0 '***' 0.001 '**' 0.01 '*' 0.05 '.' 0.1 ' ' 1
## Permutation: free
## Number of permutations: 999
```

```
# PCoA presence/absence

PcoA_RarZOTUspa <- cmdscale(distRarZOTUspa, k = (50), eig = TRUE)
```

```
## Warning in cmdscale(distRarZOTUspa, k = (50), eig = TRUE): only 29 of the first
## 50 eigenvalues are > 0
```

```
PC_RarZOTUspa <- data.frame(scores(PcoA_RarZOTUspa))
PC_RarZOTUspa$Sample <- rownames(PC_RarZOTUspa)

PC_RarZOTUspa <- PC_RarZOTUspa %>%
    as_data_frame() %>%
    inner_join(metadata, by = "Sample")
# head(PC_RarZOTUspa)

## envfit

envfit(PC_RarZOTUspa[, 1:2] ~ as.factor(PC_RarZOTUspa$Local))
```

```
## 
## ***FACTORS:
## 
## Centroids:
##                                                    Dim1    Dim2
## as.factor(PC_RarZOTUspa$Local)Bela_Vista_River1  0.0820  0.3894
## as.factor(PC_RarZOTUspa$Local)Bela_Vista_River2  0.4406 -0.1968
## as.factor(PC_RarZOTUspa$Local)Brasilia_stream    0.2655 -0.0440
## as.factor(PC_RarZOTUspa$Local)Lake              -0.0562 -0.1436
## as.factor(PC_RarZOTUspa$Local)Parana_River       0.1781  0.2861
## as.factor(PC_RarZOTUspa$Local)Parana_River2     -0.3082 -0.0015
## as.factor(PC_RarZOTUspa$Local)Reservoir         -0.2728 -0.0729
## 
## Goodness of fit:
##                                    r2 Pr(>r)    
## as.factor(PC_RarZOTUspa$Local) 0.7965  0.001 ***
## ---
## Signif. codes:  0 '***' 0.001 '**' 0.01 '*' 0.05 '.' 0.1 ' ' 1
## Permutation: free
## Number of permutations: 999
```

```
envfit(PC_RarZOTUspa[, 1:2] ~ as.factor(PC_RarZOTUspa$Year))
```

```
## 
## ***FACTORS:
## 
## Centroids:
##                                          Dim1    Dim2
## as.factor(PC_RarZOTUspa$Year)Nineteen  0.1517  0.0283
## as.factor(PC_RarZOTUspa$Year)Twenty   -0.3034 -0.0566
## 
## Goodness of fit:
##                                   r2 Pr(>r)    
## as.factor(PC_RarZOTUspa$Year) 0.3882  0.001 ***
## ---
## Signif. codes:  0 '***' 0.001 '**' 0.01 '*' 0.05 '.' 0.1 ' ' 1
## Permutation: free
## Number of permutations: 999
```

```
# rarefy down to lowest sample
comRarOTUs <- rrarefy(t(OTUs_M), sample = min(colSums(OTUs_M)))

distRarOTUs <- vegdist(comRarOTUs, method = "bray")
distRarOTUspa <- vegdist(comRarOTUs, method = "bray", binary = TRUE)

# PCoA abu

PcoA_RarOTUs <- cmdscale(distRarOTUs, k = (50), eig = TRUE)
```

```
## Warning in cmdscale(distRarOTUs, k = (50), eig = TRUE): only 35 of the first 50
## eigenvalues are > 0
```

```
PC_RarOTUs <- data.frame(scores(PcoA_RarOTUs))
PC_RarOTUs$Sample <- rownames(PC_RarOTUs)

PC_RarOTUs <- PC_RarOTUs %>%
    as_data_frame() %>%
    inner_join(metadata, by = "Sample")
# head(PC_RarOTUs)

## envfit

envfit(PC_RarOTUs[, 1:2] ~ as.factor(PC_RarOTUs$Local))
```

```
## 
## ***FACTORS:
## 
## Centroids:
##                                                 Dim1    Dim2
## as.factor(PC_RarOTUs$Local)Bela_Vista_River1  0.1846  0.3016
## as.factor(PC_RarOTUs$Local)Bela_Vista_River2  0.2997 -0.3380
## as.factor(PC_RarOTUs$Local)Brasilia_stream    0.2350 -0.2675
## as.factor(PC_RarOTUs$Local)Lake              -0.1567 -0.1229
## as.factor(PC_RarOTUs$Local)Parana_River       0.2591  0.1900
## as.factor(PC_RarOTUs$Local)Parana_River2     -0.1560  0.1411
## as.factor(PC_RarOTUs$Local)Reservoir         -0.2545  0.1093
## 
## Goodness of fit:
##                                 r2 Pr(>r)    
## as.factor(PC_RarOTUs$Local) 0.6791  0.001 ***
## ---
## Signif. codes:  0 '***' 0.001 '**' 0.01 '*' 0.05 '.' 0.1 ' ' 1
## Permutation: free
## Number of permutations: 999
```

```
envfit(PC_RarOTUs[, 1:2] ~ as.factor(PC_RarOTUs$Year))
```

```
## 
## ***FACTORS:
## 
## Centroids:
##                                       Dim1    Dim2
## as.factor(PC_RarOTUs$Year)Nineteen  0.1705  0.0121
## as.factor(PC_RarOTUs$Year)Twenty   -0.3410 -0.0243
## 
## Goodness of fit:
##                               r2 Pr(>r)    
## as.factor(PC_RarOTUs$Year) 0.427  0.001 ***
## ---
## Signif. codes:  0 '***' 0.001 '**' 0.01 '*' 0.05 '.' 0.1 ' ' 1
## Permutation: free
## Number of permutations: 999
```

```
# PCoA presence/absence

PcoA_RarOTUspa <- cmdscale(distRarOTUspa, k = (50), eig = TRUE)
```

```
## Warning in cmdscale(distRarOTUspa, k = (50), eig = TRUE): only 26 of the first
## 50 eigenvalues are > 0
```

```
PC_RarOTUspa <- data.frame(scores(PcoA_RarOTUspa))
PC_RarOTUspa$Sample <- rownames(PC_RarOTUspa)

PC_RarOTUspa <- PC_RarOTUspa %>%
    as_data_frame() %>%
    inner_join(metadata, by = "Sample")
# head(PC_RarOTUspa)

## envfit

envfit(PC_RarOTUspa[, 1:2] ~ as.factor(PC_RarOTUspa$Local))
```

```
## 
## ***FACTORS:
## 
## Centroids:
##                                                   Dim1    Dim2
## as.factor(PC_RarOTUspa$Local)Bela_Vista_River1  0.1028 -0.3727
## as.factor(PC_RarOTUspa$Local)Bela_Vista_River2  0.3882  0.1833
## as.factor(PC_RarOTUspa$Local)Brasilia_stream    0.2503  0.0811
## as.factor(PC_RarOTUspa$Local)Lake              -0.0406  0.1808
## as.factor(PC_RarOTUspa$Local)Parana_River       0.1696 -0.2441
## as.factor(PC_RarOTUspa$Local)Parana_River2     -0.2616 -0.0498
## as.factor(PC_RarOTUspa$Local)Reservoir         -0.2841  0.0203
## 
## Goodness of fit:
##                                   r2 Pr(>r)    
## as.factor(PC_RarOTUspa$Local) 0.7859  0.001 ***
## ---
## Signif. codes:  0 '***' 0.001 '**' 0.01 '*' 0.05 '.' 0.1 ' ' 1
## Permutation: free
## Number of permutations: 999
```

```
envfit(PC_RarOTUspa[, 1:2] ~ as.factor(PC_RarOTUspa$Year))
```

```
## 
## ***FACTORS:
## 
## Centroids:
##                                         Dim1    Dim2
## as.factor(PC_RarOTUspa$Year)Nineteen  0.1504 -0.0285
## as.factor(PC_RarOTUspa$Year)Twenty   -0.3009  0.0570
## 
## Goodness of fit:
##                                 r2 Pr(>r)    
## as.factor(PC_RarOTUspa$Year) 0.418  0.001 ***
## ---
## Signif. codes:  0 '***' 0.001 '**' 0.01 '*' 0.05 '.' 0.1 ' ' 1
## Permutation: free
## Number of permutations: 999
```

```
# Abus rarefied

figPCoARarASVs <- ggplot(PC_RarASVs) + aes(x = Dim1, y = Dim2, color = Group, shape = Group,
    size = 3) + xlab("PCoA1") + ylab("PCoA2") + scale_fill_manual(values = cols) +
    scale_color_manual(values = cols) + scale_shape_manual(values = forms) + ggtitle("A) ASVs abundance") +
    geom_point(position = position_jitter(0.1)) + theme(panel.grid.major = element_blank(),
    panel.grid.minor = element_blank(), panel.background = element_blank(), plot.title = element_text(size = 15L),
    plot.caption = element_text(size = 12L), axis.line = element_line(colour = "black"),
    panel.border = element_rect(colour = "black", fill = NA, size = 1))

figPCoARarZOTUs <- ggplot(PC_RarZOTUs) + aes(x = Dim1, y = Dim2, color = Group, shape = Group,
    size = 3) + xlab("PCoA1") + ylab("PCoA2") + scale_fill_manual(values = cols) +
    scale_color_manual(values = cols) + scale_shape_manual(values = forms) + ggtitle("C) ZOTUs abundance") +
    geom_point(position = position_jitter(0.1)) + theme(panel.grid.major = element_blank(),
    panel.grid.minor = element_blank(), panel.background = element_blank(), plot.title = element_text(size = 15L),
    plot.caption = element_text(size = 12L), axis.line = element_line(colour = "black"),
    panel.border = element_rect(colour = "black", fill = NA, size = 1))

figPCoARarOTUs <- ggplot(PC_RarOTUs) + aes(x = Dim1, y = Dim2, color = Group, shape = Group,
    size = 3) + xlab("PCoA1") + ylab("PCoA2") + scale_fill_manual(values = cols) +
    scale_color_manual(values = cols) + scale_shape_manual(values = forms) + ggtitle("E) OTUs abundance") +
    geom_point(position = position_jitter(0.1)) + theme(panel.grid.major = element_blank(),
    panel.grid.minor = element_blank(), panel.background = element_blank(), plot.title = element_text(size = 15L),
    plot.caption = element_text(size = 12L), axis.line = element_line(colour = "black"),
    panel.border = element_rect(colour = "black", fill = NA, size = 1))

# presence/absence rarefaction

figPCoARarASVspa <- ggplot(PC_RarASVspa) + aes(x = Dim1, y = Dim2, color = Group,
    shape = Group, size = 3) + xlab("PCoA1") + ylab("PCoA2") + scale_fill_manual(values = cols) +
    scale_color_manual(values = cols) + scale_shape_manual(values = forms) + ggtitle("B) ASVs presence/absence") +
    geom_point(position = position_jitter(0.1)) + theme(panel.grid.major = element_blank(),
    panel.grid.minor = element_blank(), panel.background = element_blank(), plot.title = element_text(size = 15L),
    plot.caption = element_text(size = 12L), axis.line = element_line(colour = "black"),
    panel.border = element_rect(colour = "black", fill = NA, size = 1))

figPCoARarZOTUspa <- ggplot(PC_RarZOTUspa) + aes(x = Dim1, y = Dim2, color = Group,
    shape = Group, size = 3) + xlab("PCoA1") + ylab("PCoA2") + scale_fill_manual(values = cols) +
    scale_color_manual(values = cols) + scale_shape_manual(values = forms) + ggtitle("D) ZOTUs presence/absence") +
    geom_point(position = position_jitter(0.1)) + theme(panel.grid.major = element_blank(),
    panel.grid.minor = element_blank(), panel.background = element_blank(), plot.title = element_text(size = 15L),
    plot.caption = element_text(size = 12L), axis.line = element_line(colour = "black"),
    panel.border = element_rect(colour = "black", fill = NA, size = 1))

figPCoARarOTUspa <- ggplot(PC_RarOTUspa) + aes(x = Dim1, y = Dim2, color = Group,
    shape = Group, size = 3) + xlab("PCoA1") + ylab("PCoA2") + scale_fill_manual(values = cols) +
    scale_color_manual(values = cols) + scale_shape_manual(values = forms) + ggtitle("F) OTUs presence/absence") +
    geom_point(position = position_jitter(0.1)) + theme(panel.grid.major = element_blank(),
    panel.grid.minor = element_blank(), panel.background = element_blank(), plot.title = element_text(size = 15L),
    plot.caption = element_text(size = 12L), axis.line = element_line(colour = "black"),
    panel.border = element_rect(colour = "black", fill = NA, size = 1))

# Plot all rarefied

figPCoARarASVs$data$Group <- as.character(figPCoARarASVs$data$Group)
figPCoARarASVs$data$Group <- factor(figPCoARarASVs$data$Group, levels = newSOrder)

figPCoARarZOTUs$data$Group <- as.character(figPCoARarZOTUs$data$Group)
figPCoARarZOTUs$data$Group <- factor(figPCoARarZOTUs$data$Group, levels = newSOrder)

figPCoARarOTUs$data$Group <- as.character(figPCoARarOTUs$data$Group)
figPCoARarOTUs$data$Group <- factor(figPCoARarOTUs$data$Group, levels = newSOrder)

figPCoARarASVspa$data$Group <- as.character(figPCoARarASVspa$data$Group)
figPCoARarASVspa$data$Group <- factor(figPCoARarASVspa$data$Group, levels = newSOrder)

figPCoARarZOTUspa$data$Group <- as.character(figPCoARarZOTUspa$data$Group)
figPCoARarZOTUspa$data$Group <- factor(figPCoARarZOTUspa$data$Group, levels = newSOrder)

figPCoARarOTUspa$data$Group <- as.character(figPCoARarOTUspa$data$Group)
figPCoARarOTUspa$data$Group <- factor(figPCoARarOTUspa$data$Group, levels = newSOrder)


figPCoARar <- ggarrange(figPCoARarASVs, figPCoARarASVspa, figPCoARarZOTUs, figPCoARarZOTUspa,
    figPCoARarOTUs, figPCoARarOTUspa, common.legend = TRUE, legend = "bottom", nrow = 3,
    ncol = 2)
ggsave("figPCoARar.tiff", figPCoARar, width = 8, height = 12, device = tiff)
figPCoARar
```

```
# Read tables

Res <- read.csv("Res_trad.csv")
Channel <- read.csv("Channel_trad.csv")
Mouth <- read.csv("Foz_trad.csv")

# Merge tables

Trad1 <- full_join(Mouth, Channel, by = "Species")
Trad <- full_join(Trad1, Res, by = "Species")
nrow(Trad)
```

```
## [1] 218
```

```
# Replace NA for zeros

Trad[is.na(Trad)] <- 0

#### transform traditional table to matrix

Trad_M <- as.matrix(Trad[, -1])  #here we remove the name column
# head(ASVs_M)

dimnames(Trad_M) <- list(Trad[, 1], colnames(Trad)[-1])  #here we name the rows

Trad_M <- Trad_M[which(rowSums(Trad_M) > 0), ]
Trad_M <- Trad_M[, which(colSums(Trad_M) > 0)]
nrow(Trad_M)
```

```
## [1] 138
```

```
# Read traditional metadata table

metadataTrad <- read.csv("metadata_trad.csv")
row.names(metadataTrad) <- metadataTrad$Sample  #here we name the rows
nrow(metadataTrad)
```

```
## [1] 39
```

```
# head(metadataTrad)
```

```
##### Diversity Estimates########

Chao1 <- data.frame(estimateR(t(Trad_M)))
Chao1t <- t(Chao1)

Shannon <- data.frame(diversity(t(Trad_M), index = "shannon"))


alphadiv <- cbind(Chao1t, Shannon)

names(alphadiv) <- c("Observed", "Chao1", "se.Chao1", "ACE", "se.ACE", "Shannon")

# head(alphadiv)

alpha_met <- merge(alphadiv, metadataTrad, by = "row.names")

# write.csv(alpha_met, 'alpha_met_Trad1.csv')

alpha_met_Trad <- read.csv("alpha_met_Trad.csv")

colsTrad <- c("#c6dbef", "#fcbba1", "#c7e9c0", "#9ecae1", "#fc9272", "#a1d99b", "#6baed6",
    "#fb6a4a", "#74c476", "#3182bd", "#de2d26", "#31a354", "#08519c", "#006d2c")

# Figure

Fig_Div_Trad <- ggplot(alpha_met_Trad) + aes(x = Group, y = Alpha, colour = Group,
    shape = Local) + geom_point(size = 3) + scale_color_manual(values = colsTrad) +
    scale_shape_manual(values = c(16, 15, 17)) + theme(axis.text.x = element_blank(),
    axis.ticks.x = element_blank(), panel.grid.major.x = element_line(size = 0.5,
        colour = "#f0f0f0"), panel.grid.major.y = element_blank(), panel.grid.minor = element_blank(),
    panel.background = element_blank(), axis.line = element_line(colour = "black"),
    panel.border = element_rect(colour = "black", fill = NA, size = 1), legend.position = ("botton")) +
    facet_wrap(vars(Measure))

newSOrder2 = c("Mouth_Seventeen", "Channel_Seventeen", "Reservoir_Seventeen", "Mouth_Eighteen",
    "Channel_Eighteen", "Reservoir_Eighteen", "Mouth_Nineteen", "Channel_Nineteen",
    "Reservoir_Nineteen", "Mouth_Twenty", "Channel_Twenty", "Reservoir_Twenty", "Mouth_TwentyOne",
    "Reservoir_TwentyOne")

Fig_Div_Trad$data$Group <- as.character(Fig_Div_Trad$data$Group)
Fig_Div_Trad$data$Group <- factor(Fig_Div_Trad$data$Group, levels = newSOrder2)

ggsave("fig_Alpha_Diversity.tiff", plot = Fig_Div_Trad, width = 9, height = 6)

Fig_Div_Trad
```

```
# rarefy down to lowest sample

distTrad <- vegdist(t(Trad_M), method = "bray")
disTradpa <- vegdist(t(Trad_M), method = "bray", binary = TRUE)

# PCoA abu

PcoA_Trad <- cmdscale(distTrad, k = 35, eig = TRUE)
```

```
## Warning in cmdscale(distTrad, k = 35, eig = TRUE): only 29 of the first 35
## eigenvalues are > 0
```

```
Pc_Trad <- data.frame(scores(PcoA_Trad))
Pc_Trad$Sample <- rownames(Pc_Trad)

Pc_Trad <- Pc_Trad %>%
    as_data_frame() %>%
    inner_join(metadataTrad, by = "Sample")
# head(PC_Trad)

## envfit

envfit(Pc_Trad[, 1:2] ~ as.factor(Pc_Trad$Local))
```

```
## 
## ***FACTORS:
## 
## Centroids:
##                                      Dim1    Dim2
## as.factor(Pc_Trad$Local)Channel    0.2276 -0.0608
## as.factor(Pc_Trad$Local)Mouth     -0.2189 -0.1617
## as.factor(Pc_Trad$Local)Reservoir -0.0249  0.1929
## 
## Goodness of fit:
##                              r2 Pr(>r)    
## as.factor(Pc_Trad$Local) 0.5979  0.001 ***
## ---
## Signif. codes:  0 '***' 0.001 '**' 0.01 '*' 0.05 '.' 0.1 ' ' 1
## Permutation: free
## Number of permutations: 999
```

```
envfit(Pc_Trad[, 1:2] ~ as.factor(Pc_Trad$Year))
```

```
## 
## ***FACTORS:
## 
## Centroids:
##                                     Dim1    Dim2
## as.factor(Pc_Trad$Year)Eighteen  -0.0484  0.0589
## as.factor(Pc_Trad$Year)Nineteen   0.1078  0.0326
## as.factor(Pc_Trad$Year)Seventeen  0.0414 -0.1465
## as.factor(Pc_Trad$Year)Twenty     0.0737  0.0723
## as.factor(Pc_Trad$Year)TwentyOne -0.1550 -0.0315
## 
## Goodness of fit:
##                             r2 Pr(>r)
## as.factor(Pc_Trad$Year) 0.1373  0.279
## Permutation: free
## Number of permutations: 999
```

```
# PCoA presence/absence

PcoA_Tradpa <- cmdscale(disTradpa, k = 35, eig = TRUE)
```

```
## Warning in cmdscale(disTradpa, k = 35, eig = TRUE): only 24 of the first 35
## eigenvalues are > 0
```

```
PC_Tradpa <- data.frame(scores(PcoA_Tradpa))
PC_Tradpa$Sample <- rownames(PC_Tradpa)

PC_Tradpa <- PC_Tradpa %>%
    as_data_frame() %>%
    inner_join(metadataTrad, by = "Sample")
# head(PC_RarOTUspa)

## envfit

envfit(PC_Tradpa[, 1:2] ~ as.factor(PC_Tradpa$Local))
```

```
## 
## ***FACTORS:
## 
## Centroids:
##                                        Dim1    Dim2
## as.factor(PC_Tradpa$Local)Channel    0.2145  0.1575
## as.factor(PC_Tradpa$Local)Mouth     -0.2832  0.0816
## as.factor(PC_Tradpa$Local)Reservoir  0.0417 -0.2144
## 
## Goodness of fit:
##                                r2 Pr(>r)    
## as.factor(PC_Tradpa$Local) 0.8733  0.001 ***
## ---
## Signif. codes:  0 '***' 0.001 '**' 0.01 '*' 0.05 '.' 0.1 ' ' 1
## Permutation: free
## Number of permutations: 999
```

```
envfit(PC_Tradpa[, 1:2] ~ as.factor(PC_Tradpa$Year))
```

```
## 
## ***FACTORS:
## 
## Centroids:
##                                       Dim1    Dim2
## as.factor(PC_Tradpa$Year)Eighteen  -0.0099  0.0235
## as.factor(PC_Tradpa$Year)Nineteen   0.1142 -0.0302
## as.factor(PC_Tradpa$Year)Seventeen -0.0228  0.0123
## as.factor(PC_Tradpa$Year)Twenty    -0.0225 -0.0731
## as.factor(PC_Tradpa$Year)TwentyOne -0.0652 -0.0696
## 
## Goodness of fit:
##                               r2 Pr(>r)
## as.factor(PC_Tradpa$Year) 0.0441  0.934
## Permutation: free
## Number of permutations: 999
```

```
# Abus rarefied

figPCoATrad <- ggplot(Pc_Trad) + aes(x = Dim1, y = Dim2, color = Group, shape = Local,
    size = 3) + xlab("PCoA1") + ylab("PCoA2") + scale_fill_manual(values = colsTrad) +
    scale_color_manual(values = colsTrad) + scale_shape_manual(values = c(16, 15,
    17)) + ggtitle("A) Fish's species abundance") + geom_point(position = position_jitter(0.1)) +
    theme(panel.grid.major = element_blank(), panel.grid.minor = element_blank(),
        panel.background = element_blank(), plot.title = element_text(size = 15L),
        plot.caption = element_text(size = 12L), axis.line = element_line(colour = "black"),
        panel.border = element_rect(colour = "black", fill = NA, size = 1))

# presence/absence rarefaction

figPCoATradpa <- ggplot(PC_Tradpa) + aes(x = Dim1, y = Dim2, color = Group, shape = Local,
    size = 3) + xlab("PCoA1") + ylab("PCoA2") + scale_fill_manual(values = colsTrad) +
    scale_color_manual(values = colsTrad) + scale_shape_manual(values = c(16, 15,
    17)) + ggtitle("B) Fish's species presence/absence") + geom_point(position = position_jitter(0.1)) +
    theme(panel.grid.major = element_blank(), panel.grid.minor = element_blank(),
        panel.background = element_blank(), plot.title = element_text(size = 15L),
        plot.caption = element_text(size = 12L), axis.line = element_line(colour = "black"),
        panel.border = element_rect(colour = "black", fill = NA, size = 1))

# Plot
figPCoATrad$data$Group <- as.character(figPCoATrad$data$Group)
figPCoATrad$data$Group <- factor(figPCoATrad$data$Group, levels = newSOrder2)

figPCoATradpa$data$Group <- as.character(figPCoATradpa$data$Group)
figPCoATradpa$data$Group <- factor(figPCoATradpa$data$Group, levels = newSOrder2)


figPCoATrad <- ggarrange(figPCoATrad, figPCoATradpa, common.legend = TRUE, legend = "bottom",
    ncol = 2)
ggsave("figPCoATrad.tiff", plot = figPCoATrad, width = 8, height = 8)
figPCoATrad
```
